# Tissue-Specific Experimental Evolution Reveals Adaptive Trade-Offs in the Plant Vascular Pathogen *Clavibacter michiganensis*

**DOI:** 10.1101/2025.10.05.680514

**Authors:** Raj Kumar Verma, Veronica Roman-Reyna, Nathan Benmoche, Evanson Ngugi, Eduard Belausov, Maya Bar, Jonathan M. Jacobs, Doron Teper

## Abstract

The plant pathogenic bacterium *Clavibacter michiganensis* (Cm) is a systemic vascular pathogen that colonizes both xylem vessels and the intracellular apoplast during different stages of infection. To identify traits and loci associated with adaptation to these distinct host microenvironments, we conducted tissue-specific experimental evolution. Twenty independent Cm lineages were repeatedly passaged in either tomato stems or leaves to promote adaptation to vascular or apoplastic lifestyles, respectively. After fifteen passages, adapted clones were characterized for virulence and virulence-related traits. These characterizations demonstrated clear differential associations of virulence-associated traits with the adapted tissue. The majority of vascular-adapted clones displayed enhanced surface attachment, reduced cellulase activity, reduced exopolysaccharide (EPS) production, and attenuated virulence on tomato compared to the parent clone. On the other hand, apoplast-adapted clones displayed reduced biofilm formation and enhanced EPS production while maintaining their virulence on tomato. Whole-genome sequencing of all adapted clones revealed candidate loci linked to tissue adaptation. Notably, six of ten vascular-adapted clones carried two independent mutations in *CMM_1284*, a putative HipB/XRE-type transcriptional regulator. A *CMM_1284* marker exchange mutant displayed phenotypes similar to vascular-adapted clones, suggesting a role for this regulator in vascular colonization. Together, these findings highlight the role of phenotypic plasticity in tissue adaptation of plant pathogens, showing that tissue-specific adaptation involves modulation of surface attachment, EPS production, and cell wall–degrading enzymes. They further reveal a regulated trade-off between vascular persistence, supported by strong surface attachment, and systemic virulence, which depends on bacterial dispersal and migration.

## Introduction

Plant pathogenic microorganisms specialize in colonizing host tissues according to lifestyle: systemic pathogens spread through the vascular system, while non-systemic ones remain confined to apoplastic regions. These contrasting lifestyles shape disease etiology, epidemiology, and management. Non-systemic bacteria colonize the apoplast, inducing galls or necrotic lesions [1–3], spread locally by water splash, wounding, or insects, and rarely invade the vasculature [4]. Vascular pathogens specialize in xylem or phloem colonization. Phloem-associated pathogens are insect-transmitted, phloem-restricted, passively transported, and have reduced, low-GC genomes [5]. Xylem-associated pathogens show diverse transmission and tissue specificity: *Xylella* spp. are strictly xylem-limited, whereas vascular Xanthomonads, *Ralstonia* spp., *Erwinia amylovora*, and *Clavibacter* spp. also colonize apoplast tissues [6,7]. In *E. amylovora*, transitions between xylem and parenchyma are central: it enters the xylem from floral parenchyma, spreads systemically, and exits into bark parenchyma, causing tissue degradation and proliferation [8]. Mechanisms of tissue specificity and lifestyle transitions remain unclear; even related strains differ in tropism. In *Xanthomonas*, the glycosyl hydrolase gene *cbsA* distinguishes vascular from non-vascular lifestyles [9]. Xylem-associated bacteria produce biofilm-like matrices attaching to vessel walls, essential for systemic spread and implicated in drought-like symptoms by blocking water flow [10]. Mutational analyses show that balancing biofilm formation and dispersal is critical for systemic colonization [11,12].

Bacterial canker, caused by the Actinomycetota *Clavibacter michiganensis* (Cm), is one of the most destructive tomato diseases [13]. Symptoms include canker lesions on stems and branches, leaf wilting, necrosis, and ‘bird’s-eye’ spots on fruits [13]. Cm spreads by colonizing xylem vessels, causing vascular collapse through blockage of water-conducting elements [14]. The pathogen relies heavily on secreted hydrolases such as cellulases, xylanases, pectate lyases, and serine proteases, encoded mainly by the *chp/tomA* pathogenicity island (PAI) and plasmids pCM1 and pCM2 [15,16]. Variation in these regions strongly correlates with pathogenicity [17–19], and disruption of specific hydrolases such as *celA*, *chpC*, and *pat-1* reduces virulence [20–23]. Microscopy has shown Cm forming biofilm-like aggregates attached to xylem walls [24], suggesting biofilms aid systemic colonization, though their composition, structure, and regulation are poorly understood. Only a few studies indirectly link biofilm formation to regulatory networks such as the stringent response and cell wall integrity [25,26]. Despite evidence for surface attachment *in vitro* and *in planta*, adhesion mechanisms remain undefined and no adhesion-associated macromolecules have been identified. Cm exopolysaccharides (EPS) were hypothesized to contribute to virulence but this hypothesize have yet to be tested by functional or genetic analyses [27]. The model strain NCPPB382 harbors at least four predicted EPS gene clusters [16,28], but their role in EPS production, xylem association, and virulence is unvalidated. Cm also colonizes the leaf apoplast, producing blister-like structures [15,29], though the contribution of this lifestyle to disease progression remains unclear.

In this study, we employed a tissue-based experimental evolution approach to identify traits and genetic loci selected for adaptation to either the vascular or apoplastic lifestyle of Cm.

## Results

### Cm migrates from the xylem to apoplast tissue during infection

Cm is traditionally classified as a xylem-inhabiting bacterium [7]. However, in contrast to xylem-restricted bacteria such as *Xylella fastidiosa*, Cm has also been reported to colonize intracellular apoplastic spaces within host tissues [7,24,30]. To assess Cm’s tissue localization, GFP-labeled Cm [24] was monitored for colonization throughout the plant following infection.

First, we examined Cm colonization of the apoplast by syringe-infiltrating bacteria into tomato leaves (Fig. 1A–C). Cm caused localized water-soaked lesions that became necrotic within 5–10 days (Fig. 1A), accompanied by a ∼10,000-fold population increase by 3 days post-infiltration (dpi) (Fig. 1B). Confocal microscopy showed Cm cells in intracellular apoplastic spaces of leaf parenchyma (Fig. 1C), confirming its ability to thrive during apoplastic infection. Like many apoplastic pathogens, Cm-induced lesions stayed confined to infiltrated areas, and inoculated plants showed no wilt or canker symptoms typical of systemic infection. Thus, while Cm can act systemically, direct apoplastic introduction rarely leads to systemic spread.

**Fig. 1.**
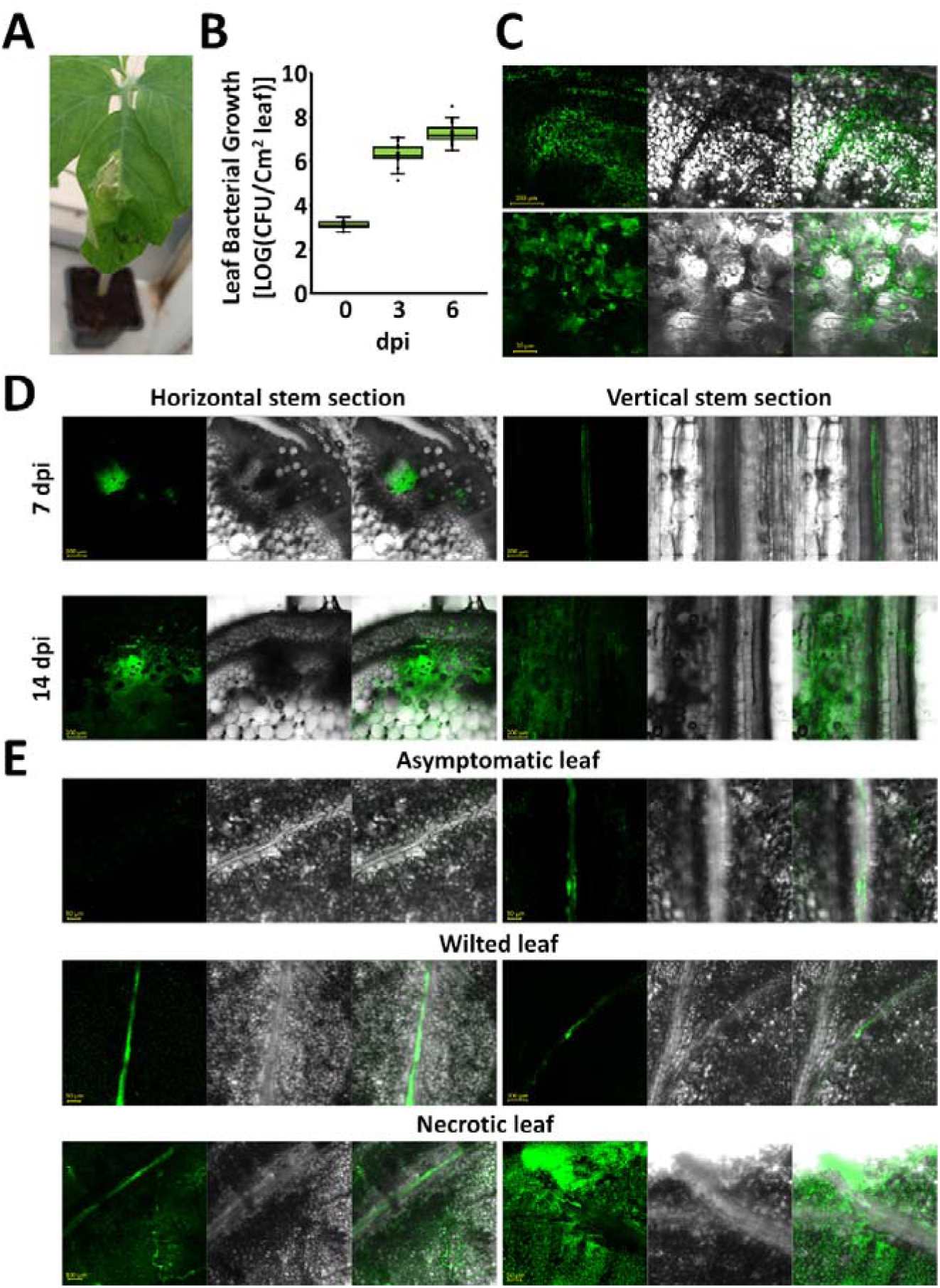
*In situ* localization of Cm bacteria during infection. (**A**–**C**). Cultures (10^4^ CFU/ml) of GFP-labeled Cm were infiltrated into six-leaf-stage tomato leaves using a needless syringe. (**A**) Representative leaf photographed 10 days post-infiltration (dpi). (**B**) Bacterial population at the infiltration site on days 0, 3, and 6 dpi. Data represent 26 biological repeats pooled from three independent experiments. (**C**) Infiltrated leaf tissues were visualized by confocal laser scanning microscopy using a GFP filter at 7 dpi. (**D**, **E**) Stem areas between the cotyledons of four-leaf-stage tomato plants were wound-inoculated using toothpicks soaked in 10^7^ CFU/ml GFP-labeled Cm bacterial cultures. Bacteria were visualized by confocal laser scanning microscopy with a GFP filter. (**D**) *In situ* localization of Cm at 3 cm above the infection site, visualized in horizontal and vertical stem cross sections of Cm-inoculated plants at 7 dpi (prior to symptom development) and 14 dpi (when symptoms are apparent). (**E**) *In situ* localization of Cm in asymptomatic, wilted, and necrotic leaves at 14 dpi. The right and left panels show the two most common localization patterns at leaves during the described symptomatic stages. All depicted data represent at least 10 biological repeats conducted in at least two (C) or three (A, B, D, E) independent experiments.

Next, we examined Cm localization during systemic infection by wound-inoculating stems with GFP-labeled Cm between the cotyledons. *In situ* monitoring was done 3 cm above the infection site at 7 dpi, before severe symptoms, and at 14 dpi, when ∼75% of leaflets were wilted and stem cankers were apparent. At 7 dpi, Cm was mainly in xylem vessels with little or no presence in pith parenchyma, consistent with its xylem-restricted lifestyle (Fig. 1D, upper panels). By 14 dpi, bacteria had burst from the xylem into pith parenchyma intracellular spaces (Fig. 1D, lower panels). In distal leaf tissues at 14 dpi, asymptomatic leaflets showed no bacteria or restriction to the main vein, wilted leaflets contained bacteria in vascular bundles or nearby tissues, and necrotic leaflets were heavily colonized in both vascular bundles and apoplastic parenchyma (Fig. 1E). Together, these findings indicate that Cm exits the vascular system into apoplastic parenchyma during late infection, and that xylem escape is key to its infection cycle.

### Experimental evolution of Cm under distinct vascular and apoplastic niches

The xylem and apoplast microenvironments within a plant host differ greatly and pose unique challenges to invading pathogens [6]. Since Cm can thrive and transition between these habitats, it likely possesses traits supporting survival in each, along with phenotypic plasticity enabling the shift. We hypothesized that traits or loci associated with each environment could be identified by experimentally evolving Cm under repeated exposure to vascular or apoplastic conditions. To test this, we generated 20 parallel tissue-adapted Cm populations by repeated passaging in vascular or apoplastic niches. Ten clones were evolved in vascular tissue (S1–S10) and ten in the leaf apoplast (L1–L10). Experiments used Cm NCPPB382, hereafter referred to as Cm WT or parent clone. Vascular-adapted clones were passaged through distal stems: tomato stems were inoculated via toothpick puncture, and bacteria were re-isolated 14–21 days later from tissues 6 cm above the inoculation site. Isolates were pooled and used to inoculate new hosts for the next passage (Fig. 2). Apoplast-adapted clones were passaged by infiltrating diluted Cm suspensions (10□ CFU/ml) into leaf apoplast with a needleless syringe. Bacteria were re-isolated 7–10 days later from the infiltration area, pooled, and used to inoculate new hosts (Fig. 2). Parallel passaging continued for 15 cycles, after which one representative clone from each lineage was selected for phenotypic and genomic analyses.

**Fig. 2.**
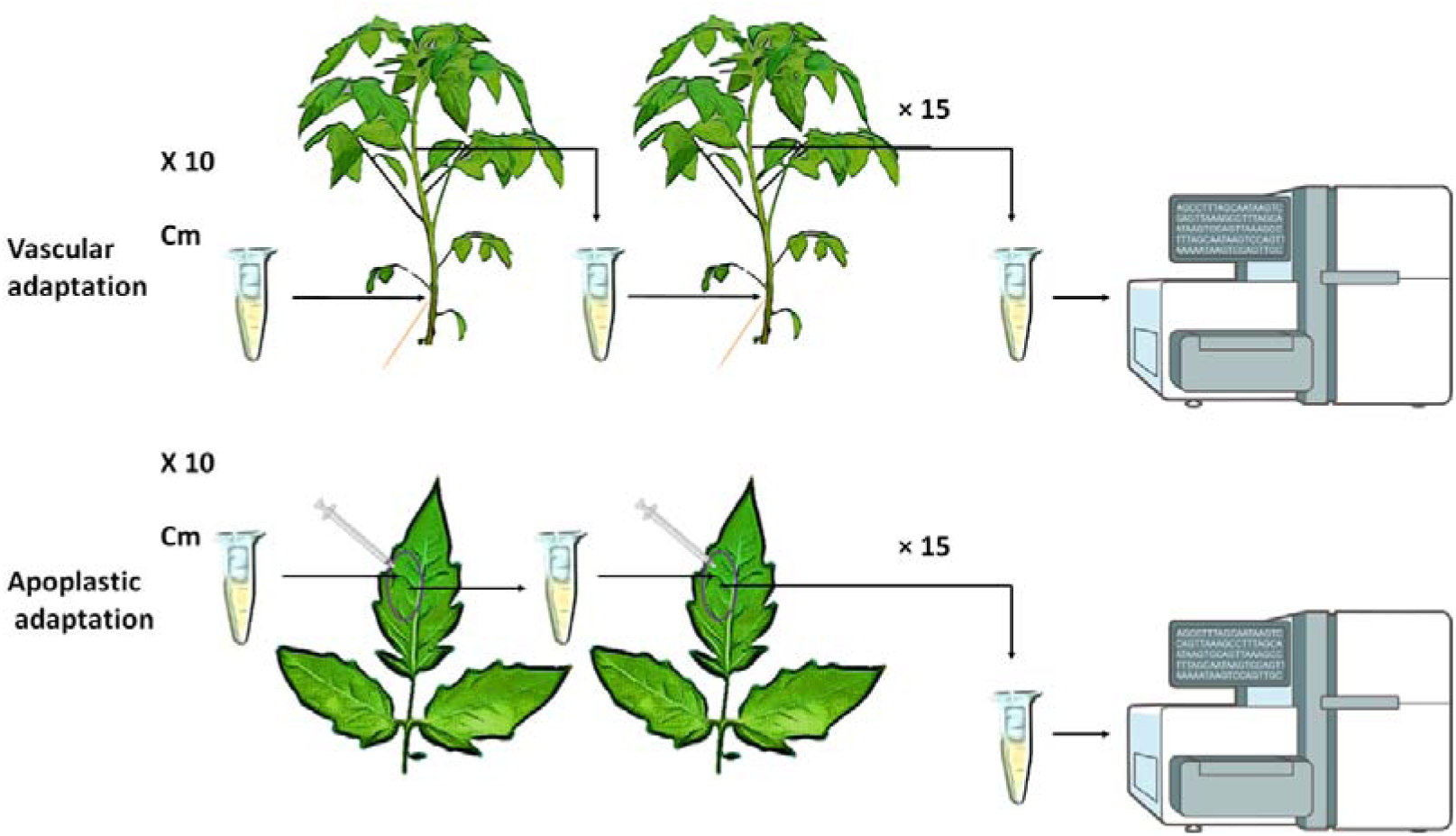
Cartoon illustrating the experimental evolution procedure used in the study. Ten parallel Cm lines underwent repeated inoculation and re-isolation passages lasting 14–21 days, performed via stem (upper panel, vascular adaptation) and leaf (lower panel, apoplastic adaptation) inoculations. The cartoon was created with the assistance of cartoon photo editor (https://play.google.com/store/apps/details?id=com.gamebrain.cartoon&hl=en) and NIAID Visual & Medical Arts (bioart.niaid.nih.gov/bioart/386, bioart.niaid.nih.gov/bioart/506).

### Vascular-adapted clones exhibit reduced vascular and apoplastic virulence

We investigated whether adaptation to specific plant tissues affected virulence and colonization in vascular and apoplastic tissues. After 15 passages, adapted clones and the parent Cm WT were inoculated into tomato stems or leaves, and plants were monitored for symptoms and bacterial growth. For vascular virulence, stems were toothpick-inoculated and wilt symptoms tracked for 21 days. Bacterial titers were also measured 1 and 10 cm above the inoculation site. While apoplast-adapted clones (L1–L10) caused wilt comparable to WT, eight vascular-adapted clones (S1, S2, S3, S4, S5, S6, S8, S9) showed reduced wilt (Fig. 3A and 3B), despite unchanged bacterial titers at 21 dpi (Fig. 3C). To test kinetic systemic spread, bacterial loads at 5 and 10 cm were measured at 7 and 14 dpi in three virulence attenuated vascular-adapted clones (S2, S4, S9). By 14 dpi, S2 and S4 colonization matched WT, while S4 showed a fivefold reduction 10 cm above the inoculation site (Fig. 3D), suggesting delayed systemic spread could explain its attenuated phenotype. For apoplast virulence, bacteria were syringe-infiltrated into leaves and symptoms scored for 10 days. Apoplast-adapted clones resembled WT (Fig. S1A, S1B). Four vascular-adapted clones (S2, S3, S4, S5) showed weaker or delayed symptoms, while S7 caused stronger, earlier lesions (Fig. S1A, S1B). However, all clones eventually reached similar apoplastic titers (Fig. S1C). Overall, vascular adaptation produced clones with attenuated virulence in both tissues, whereas apoplast adaptation preserved full virulence.

**Fig. 3.**
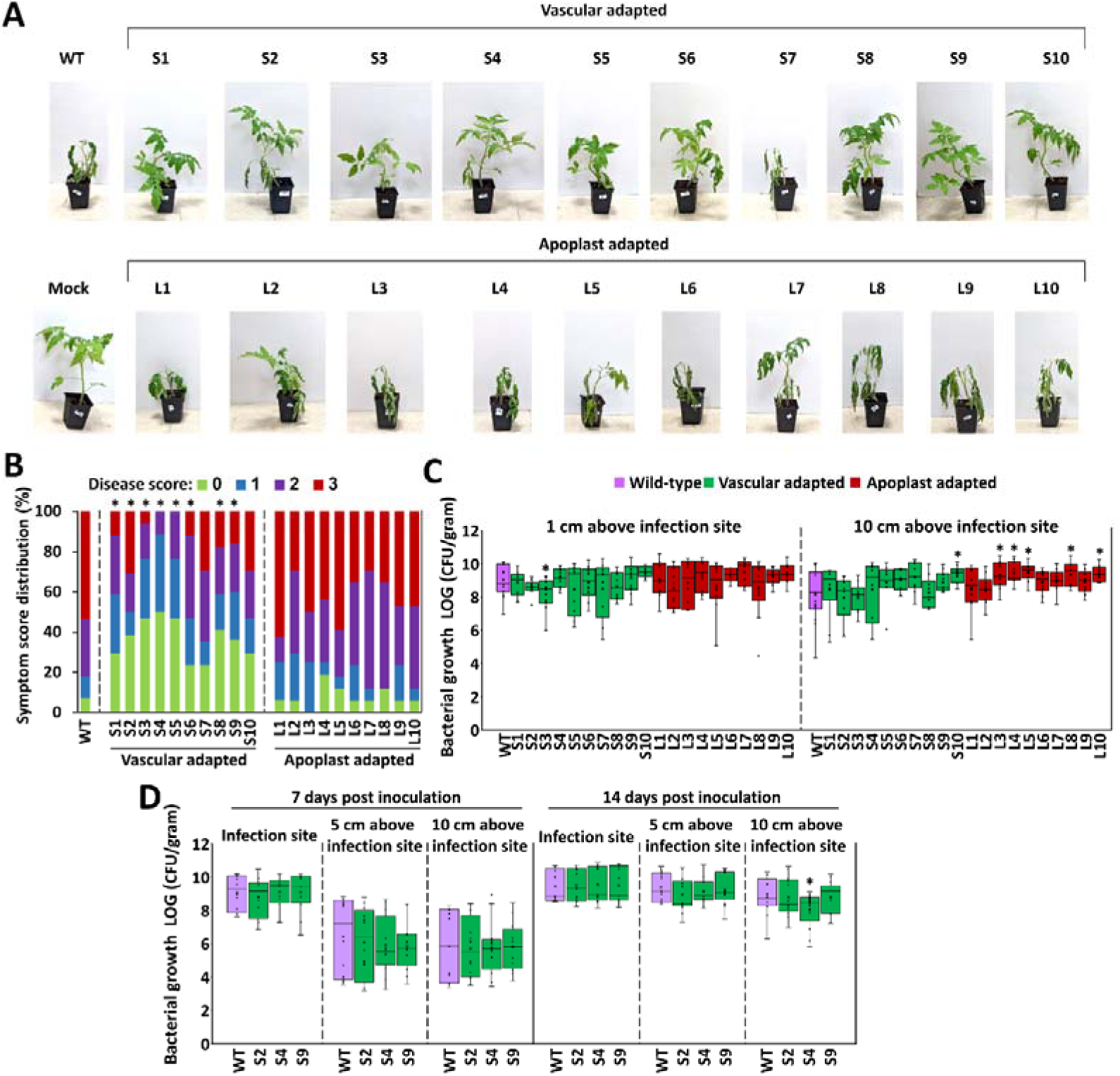
Vascular adaptations affect vascular virulence. Four-leaf stage tomato plants were wound-inoculated with the indicated vascular adapted clones (S1-S10), apoplast-adapted clones (L1-L10), and Cm382 (WT) at the stem areas between the cotyledons. **(A)** Representative plants (out of 14 repeats) were photographed at 21 dpi. (**B**) Wilt symptoms were scored by the percentage of wilted leaves according to the following scale: 0 – no wilting, 1 = 1-25%, 2= 25-50%, 3= 51-100%. The graph depicts the distribution of 14 repeats for each clone pooled from two experiments. “*” represent significant difference (chi-squared test, p-value < 0.05) compared to WT. (**C)** Bacterial growth in stems at 1 and 10 cm above the sites of infections at 21 dpi. **(D**) Bacterial growth of the indicated clones in stems at the site of infections, 5 cm above the site of infections, and 10 cm above the site infections was monitored at 7 and 14 dpi. (**C**, **D**) Data is depicted in a box plot graph (boxes represent 25%-75% of the data, line represents median) representing 10 repeats pooled from two experiments. “*” represent significant differences (U-Test, p-value < 0.05) compared to WT in the same sampling area at the same dpi.

### Vascular-adapted clones demonstrate reduced stem-to-leaf migration

Most vascular-adapted clones colonized and migrated systemically through the stem similarly to the WT parent, suggesting that their reduced wilting symptoms are not solely due to limited stem colonization. Since our *in situ* localization experiments linked wilting to bacterial presence in leaves (Fig. 1), we hypothesized that their attenuated virulence may reflect reduced migration from stem to leaves. To test this, tomato plants were stem-inoculated with Cm WT, vascular-adapted clones (S2, S4, S9), and apoplast-adapted clones (L2, L7, L9). Bacterial presence and populations were monitored in pooled terminal leaflets from the second to fourth true leaves at 7 and 14 dpi. Vascular-adapted clones showed markedly lower frequencies and titers compared to WT, whereas apoplast-adapted clones colonized leaves at equal or higher levels (Fig. 4A and 4B). At 7 dpi, over 50% of leaflets were colonized by WT and apoplast-adapted clones but only 20–30% by vascular-adapted clones. By 14 dpi, colonization in WT and apoplast-adapted clones exceeded 90%, while S2 and S4 reached ∼50% and S9 ∼75% (Fig. 4A). These results suggest vascular-adapted clones either migrate more slowly from stems or struggle to establish in leaves. To distinguish these, we directly infiltrated leaves with all seven representative clones and tracked colonization at 3, 6, and 9 dpi. No significant differences were detected (Fig. 4C), indicating reduced leaf colonization results from impaired stem-to-leaf migration rather than defective leaf colonization. We next asked if this defect reflected altered tissue localization. GFP-labeled S2 and S4 were compared with WT in distal stem and leaf tissues (Fig. S2). No clear differences in localization were observed, and patterns correlated with leaflet symptoms. Consistent with isolation data, bacteria were absent from many asymptomatic leaflets inoculated with S2 and S4, though this also occurred at lower frequency in WT.

**Fig. 4.**
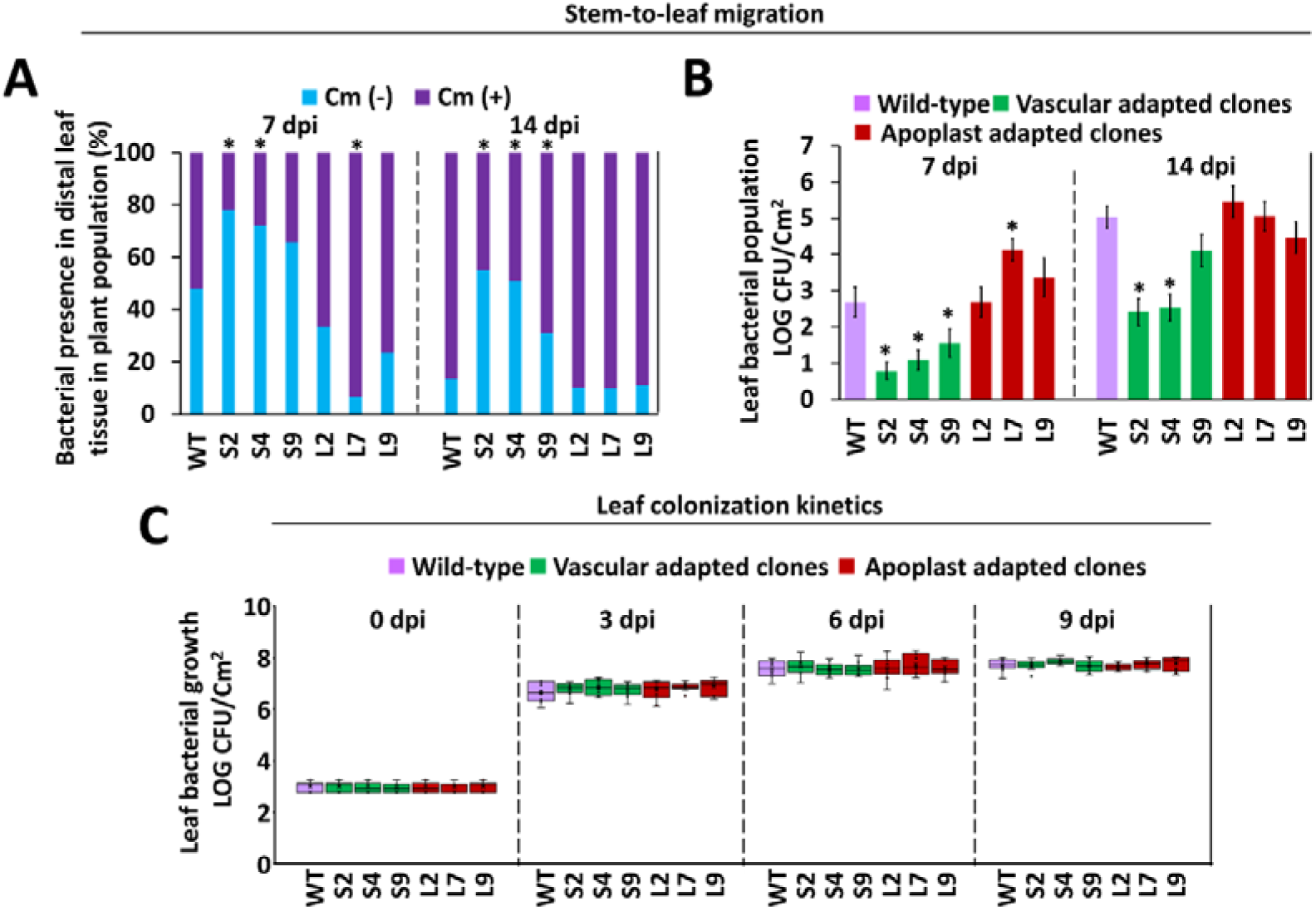
Vascular and apoplastic adaptations affect stem-to-leaf migration. (**A**, **B**) Four-leaf stage tomato plants were wound-inoculated with the indicated vascular adapted clones (S2, S4, and S9), apoplast-adapted clones (L2, L7, and L9), and Cm382 (WT) at the stem areas between the cotyledons. Bacteria were isolated and quantified from pooled samples of terminal leaflets from the second, third, and fourth true leaves of each plant at 7 and 14 days post inoculation (dpi). Data represents at least 30 repeats pooled from five (for WT, S2, S4, S9) or three (for L2, L7, and L9) experimental repeats. (**A**) Stacked bar graphs represent presence/absence of bacteria in the sampled leaflets at 7 and 14 dpi. “*” indicates that data were significantly different compared to Cm WT (chi-squared test, p-value < 0.05) at the same dpi. (**B**) Bar graphs represents the averages and standard errors of leaf bacterial populations. “*” indicates that bacterial populations were significantly different compared to Cm WT (U-test, p-value < 0.05) at the same dpi. (**C**) Six-leaf stage tomato leaves were inoculated with the indicated clones through infiltration of bacterial cultures (10^4^ CFU/ml) using a needless syringe. Bacterial growth in the infiltration sites at 0, 3, 6, and 9 dpi. Data represents 10 repeats for each clone pooled from two experiments. No significant differences (U-test) were observed between each of the clones to the WT.

In summary, vascular-adapted clones show a pronounced defect in stem-to-leaf migration, likely underlying their attenuated virulence.

### Tissue-adapted clones demonstrate altered colony morphology, production of exopolysaccharides, and surface attachment

Considering that vascular adaptation altered both vascular and non-vascular virulence, we hypothesized that traits linked to pathogenicity and xylem association were also modified. We therefore examined EPS production, surface attachment, exoenzyme activity, siderophore production, hypersensitive response (HR) elicitation in non-hosts, and expression of virulence-associated genes. Siderophore production and HR induction were unchanged in adapted clones relative to Cm WT (Fig. S3), but several traits shifted in a tissue-dependent manner.

On rich or minimal media, eight of ten vascular-adapted clones (S1, S2, S3, S4, S5, S6, S8, S9) that also showed attenuated vascular virulence formed dry, condensed colonies, whereas apoplast-adapted clones resembled WT with wet, glossy colonies (Fig. 5A). This suggested reduced EPS secretion or altered composition. Quantification confirmed seven vascular-adapted clones produced less EPS (Fig. 5B), while six apoplast-adapted clones produced significantly more. To further investigate this pattern, we examined the transcriptional expression of genes predicted to be involved in EPS production [28] in representative vascular-adapted clones (S2, S4, and S9) with low EPS yields and observed significant reductions in specific target genes in some clones, but not others (Fig. S4). Since EPS production is traditionally associated with surface attachment and biofilm formation [31,32], we next conducted *in vitro* surface attachment assays, which also serve as a proxy for biofilm formation, on all adapted clones using a 24-well crystal violet staining assay (Fig. 5C). Unexpectedly, vascular-adapted clones with dry colonies and low EPS showed stronger attachment than WT, while apoplast-adapted clones, which produced more EPS, had weaker attachment (Fig. 5C). These results suggest that, unlike in other systems, EPS may reduce attachment and biofilm formation in Cm.

**Fig. 5.**
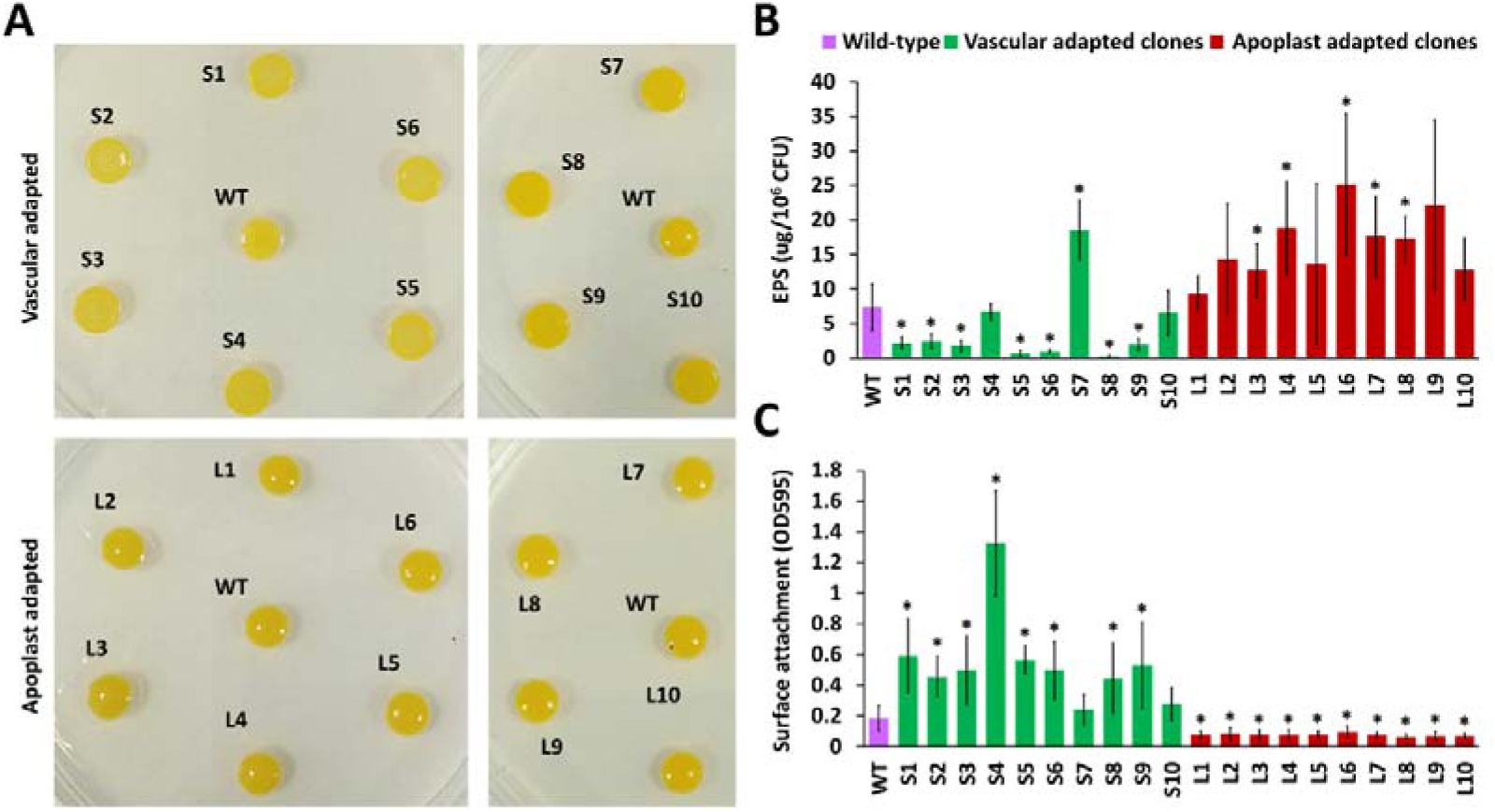
Vascular and apoplastic adaptations effect on EPS production and surface attachment. (**A**, **B**) Cm WT and the indicated vascular and apoplast adapted clones were spotted (OD600=1) on LB agar supplemented with 5% sucrose and incubated for four days. Plates were photographed at 5 dpi (**A**) and EPS was quantified in the resulting clones (**B**) using phenol–sulfuric acid hydrolysis assays following by colorimetric quantification and standardization to bacteria count. (**C)** Bacteria were grown on LB media, standardized to OD600=0.5 and incubated in 24 well plates without shaking for 10 days. Supernatants were removed, wells were washed three times with distilled water, and attached bacteria were stained with crystal violet to quantify surface attachment followed by colorimetric quantification. **(B**, **C)**. Graphs represent one out of three experimental repeats each composed of three (**B**) or four (**C**) biological repeats showcasing similar results. “*” represent significant difference (U-test, p-value < 0.05) compared to Cm WT.

Overall, EPS production and surface attachment appear shaped by tissue-specific selective pressures: vascular adaptation favored high attachment and low EPS, while apoplast adaptation favored low attachment and high EPS.

### Vascular adapted clones demonstrate reduced cellulase and amylase activity

After establishing that EPS production and surface attachment are selected during tissue adaptation, we assessed exoenzyme activity and production using plate halo assays. Cm bacteria heavily depend on secreted hydrolases such as CAZymes and serine proteases to cause disease [15]. Therefore, we monitored plate exoenzyme activity by halo assays on multiple substrates and transcriptional expression of predicted virulence associated serine proteases and CAZymes.

Halo assays revealed no tissue-adaptation bias in lipase and xylanase activity in the adapted clones (data not shown). However, a clear association was observed between tissue adaptation and amylase and cellulase activity, visualized using starch and CMC degradation, respectively (Fig. 6A and 6B). Specifically, all vascular-adapted clones demonstrated reduced amylase and cellulase activity, while no significant difference was observed in the apoplast-adapted clones compared to the WT (Fig. 6A and 6B). While amylase activity was not subjected to any study in Cm, previous studies reported that cellulase activity in Cm is mediated by the secreted endo-beta-1,4-glucanase CelA, and disruption of *celA* abolishes cellulase activity in CMC halo assays, leading to a significant reduction in virulence with minimal impact on *in planta* growth [20,33]. Therefore, we monitored the transcriptional expression of *celA* and the putative amylase encoding gene *aglC* in the representative vascular adapted clones S2, S4, and S9 by RT-qPCR, and identified that *celA* expression was reduced in all three clones while *aglC* expression was not significantly affected (Fig. 6C). To further confirm that *celA* expression is reduced in the vascular adapted clones, Cm WT, S2, S4, and S9 were introduced with a plasmid carrying transcriptional fusion of the *celA* or the *gyrB* (used as a control) promoter to a *uidA* (*GUS*) reporter gene and monitored for GUS activity. Supporting transcriptional data, *celA* promoter activity was significantly reduced in the vascular adapted clones compared to Cm WT while the *gyrB* promoter activity was not significantly altered (Fig. 6D). Next, we tested the transcriptional expression of additional CAzyme coding genes *xysA* and *nagA*, which did not change significantly compared to the Cm WT and three Chp/Pat-1 family serine proteases: *chpC* and *chpE* that were reduced in S2 and S4, and *chpG* that was unaltered (Fig. S4). These findings suggest that vascular-adaptation resulted in reduced exoenzyme activity and expression of specific secreted virulence factors.

**Fig. 6.**
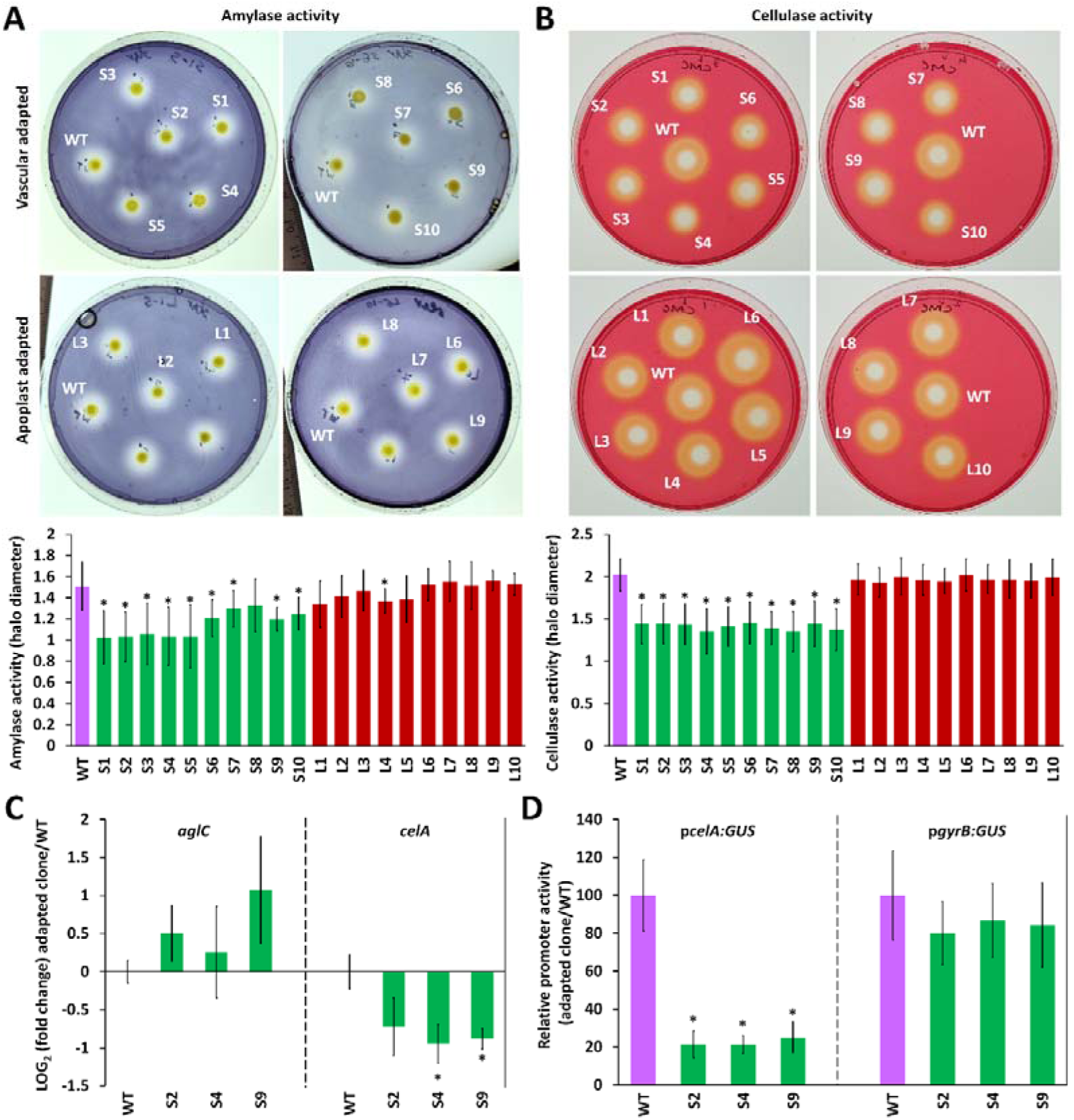
Vascular-adapted clones demonstrate reduced amylase and cellulase activity. (**A**, **B**) *Cm* WT and the indicated vascular- and apoplast-adapted clones were spotted (OD600 = 1) onto M9 medium supplemented with 0.1% starch (A) or carboxymethyl cellulose (CMC) (B), and 0.01% sucrose, then incubated for seven (A) or four (B) days. Plates were stained with 0.1% iodine (A) or Congo red (B), photographed (upper panels), and halo diameters were measured with a ruler to assess starch or CMC degradation (lower panels). Graphs represent means ± SE of at least nine replicates pooled from two independent experiments. (**C**) mRNA transcript abundance of the putative amylase-encoding gene *aglC* (CMM_2797) and the cellulase-encoding gene *celA* (pCM1_0020) was quantified by RT-qPCR in the indicated *Cm* cultures after 24 h incubation in sucrose-supplemented M9 medium. *gyrA* (CMM_0007) mRNA levels were used for normalization. Graphs represent means ± SE of relative transcript abundance compared to *Cm* WT, based on six independent biological replicates pooled from two independent experiments. (**D**) GUS activity was measured in the indicated *Cm* cultures harboring plasmids carrying *gyrB* (CMM_0006) or *celA* promoter-GUS fusions after 24 h incubation in sucrose-supplemented M9 medium. Graphs represent means ± SE of relative GUS activity compared to *Cm* WT, based on nine independent biological replicates pooled from three independent experiments. (**A**–**D**) Asterisks (*) indicate statistically significant differences compared to *Cm* WT (U-test, p-value < 0.05).

### Overexpression of *celA* in vascular-adapted clones only partially restores virulence

The cellulase CelA has been previously established as a key virulence factor in Cm that is essential for the development of wilting symptoms [20,33]. Hence, the attenuated virulence of the vascular-adapted clones may be a result of their reduced *celA* expression and, consequently, lower cellulase activity. To test this, we investigated whether ectopic expression of *celA* could restore virulence in these vascular-adapted clones. We introduced *celA*, fused to an HA epitope and driven by the *groEL* promoter from *Microbacterium esteraromaticum* (which we previously confirmed to be active in Cm, data not shown), into vascular-adapted clones S2, S4, and S9. CelA accumulation in the transformants was confirmed by restoration of cellulase activity to WT levels in plate assays and by Western blot analysis (Fig. S5A and S5B).

Next, we inoculated tomato stems with Cm WT, the vascular-adapted clones, or the *celA*-expressing vascular-adapted clones and monitored wilting symptoms. Each CelA-expressing clone displayed increased virulence relative to its parental vascular-adapted clone but remained less virulent than Cm WT (Fig. S5C). This indicates that although restoring cellulase activity partially recovers virulence, it cannot fully compensate for the reduced virulence of the vascular-adapted clones.

### Genomic analyses of vascular- and apoplastic-adapted clones

Vascular and apoplastic adaptation yielded clones with distinct phenotypes that strongly correlated to their adapted tissue. We hypothesized these changes were driven by genomic alterations and fixation of mutations underlying the observed traits. To test this, we sequenced genomes of all adapted clones used in phenotypic assays after 15 passages from each of the 20 lineages, as well as individual clones at passages 5 and 10 to assess mutation fixation over time. Three independent colonies of the parent strain Cm NCPPB382 were also sequenced for comparison. Sequencing was performed on an Illumina Nextseq2000 platform (150 bp paired-end reads) at the Applied Microbiology Services Laboratory, The Ohio State University. De novo assemblies were generated for all clones and compared with Cm NCPPB382 to detect large deletions and smaller variations via SNP and INDEL calling. Comparative analysis revealed no large deletions or plasmid loss. Mutations were limited to a small set of SNPs and INDELs (Table S1). Notably, despite major effects on cellulase activity, no mutations were detected in *celA* or its promoter in vascular-adapted clones. Tracking fixation patterns across passages 5, 10, and 15 showed some mutations arose early and persisted, while others appeared only transiently, indicating variable retention of diversity within lineages (Table S1).

Most adapted clones harbored a few mutations in putative coding genes that resulted in gene disruption or shifts in amino acid composition (Tables 1 and S1). Some mutations appeared exclusively in tissue-specific groups. A G308→A substitution in *CMM_2466*, encoding a putative restriction endonuclease, caused a G103→D change and was found in four apoplast-adapted clones (L1, L2, L3, L6) but absent in vascular clones (Table 1). Conversely, six vascular-adapted clones carried two independent mutations in *CMM_1284*, absent from apoplast clones: C58→T in S1, S2, S6 (R20C), and A110→G in S3, S4, S9 (H37R) (Table 1, Fig. S5A). CMM_1284 encodes a 107-aa HipB/XRE-type transcriptional regulator, with both substitutions located in its predicted helix-turn-helix DNA-binding domain (Fig. S6B).

**Table 1.**
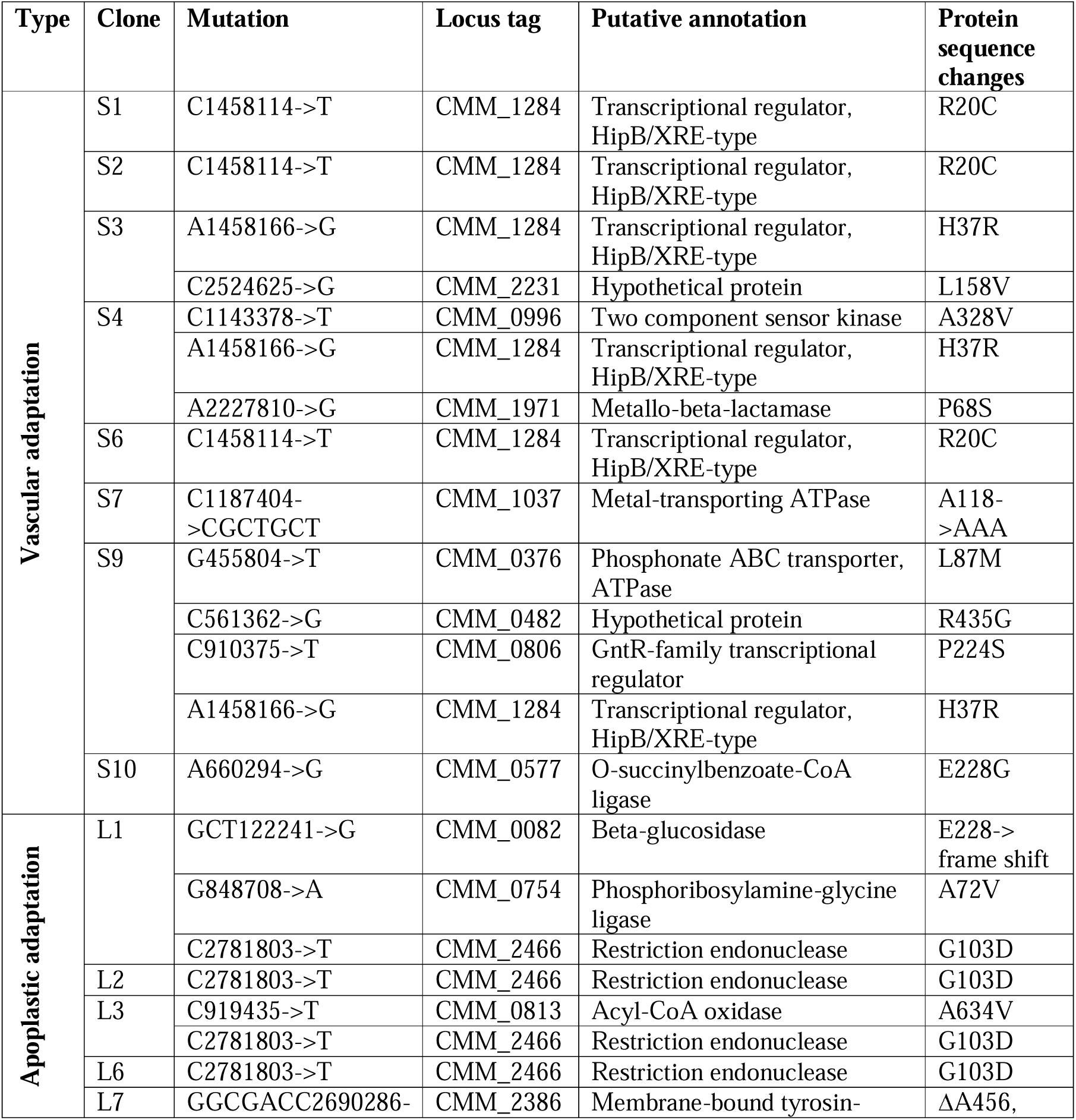
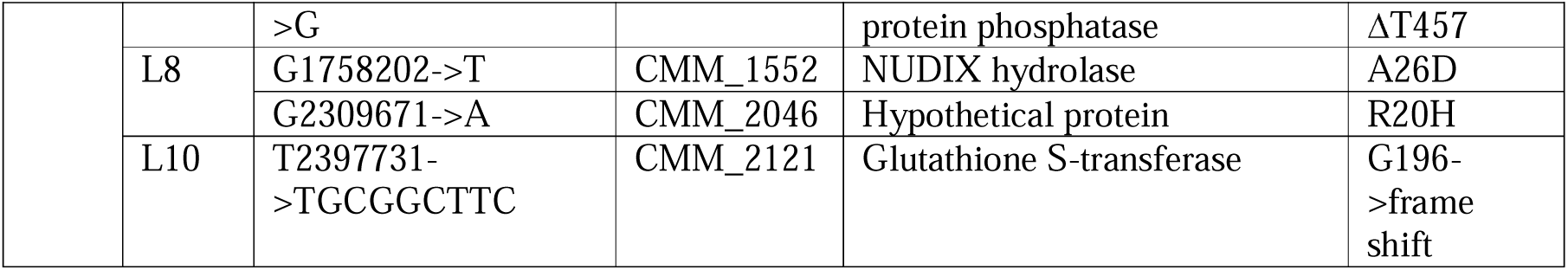
Coding-sequence substitutions and short indels in vascular and apoplast-adapted clones.

These findings reveal specific mutations selected during vascular and apoplastic adaptation, providing a genetic basis for tissue-specific traits.

### CMM_1284 modulates virulence, EPS production, surface attachment, and cellulase activity

Comparative genome analyses identified two independent mutations in the putative transcriptional regulator *gene CMM_1284* across six vascular-adapted clones, all showing attenuated virulence, enhanced surface attachment, and reduced cellulase activity. These findings suggest CMM_1284 contributes to the vascular lifestyle by modulating these traits. To test this, we constructed a *CMM_1284* marker exchange mutant (CmΩ1284; Fig. S6C and S6D) and examined its impact on traits altered in vascular-adapted clones. CmΩ1284 was first tested for virulence, systemic colonization, and stem-to-leaf migration in tomato. Like vascular-adapted clones, CmΩ1284-inoculated plants displayed significantly reduced wilting symptoms compared to WT (Fig. 7A and 7B), while systemic colonization in the stem was unaffected (Fig. 7C). In contrast, CmΩ1284 showed markedly reduced stem-to-leaf migration, with lower bacterial frequency and populations in the second to fourth leaves at 7 and 14 dpi (Fig. 7D, and 7E). Finally, we assessed CmΩ1284 for in vitro virulence traits and found that it formed dry colonies, produced less EPS, exhibited stronger surface attachment, and showed reduced cellulase activity. These features closely resemble those of most vascular-adapted clones (Fig. 7F–H).

**Fig. 7.**
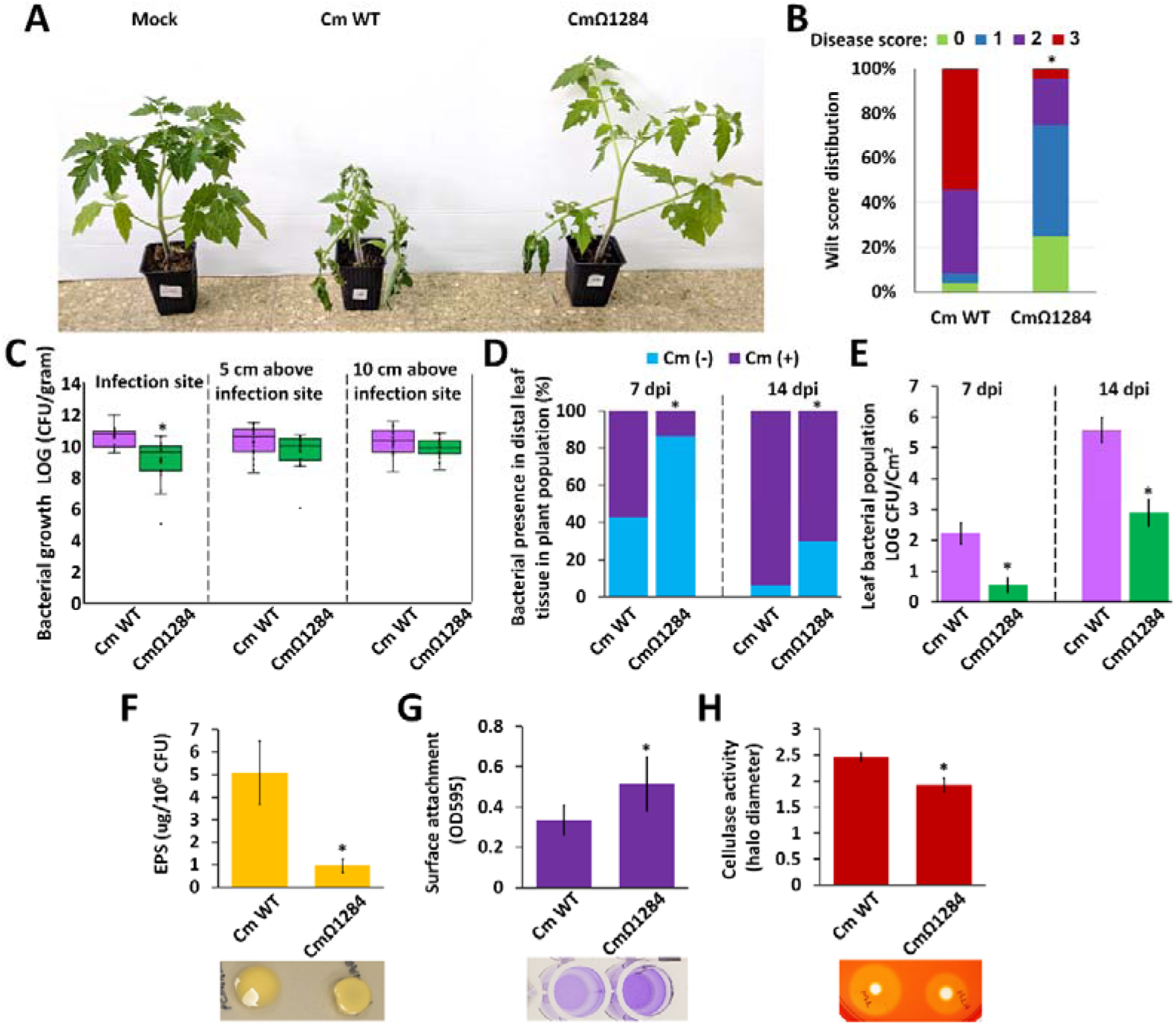
Characterization of the CMM_1284 marker exchange mutant. (**A**–**E**) Four-leaf stage tomato plants were wound-inoculated with Cm WT and the *CMM_1284* marker exchange mutant (CmΩ1284) at the stem areas between the cotyledons. (**A**) Representative plants were photographed at 14 days post-inoculation (dpi). (**B**) Wilt symptoms were scored based on the percentage of wilted leaves using the following scale: 0 = no wilting, 1 = 1–25%, 2 = 25–50%, 3 = 51–100%. The stacked graph shows the distribution of 24 replicates for each clone pooled from three experiments at 14 dpi. (**C**) Bacterial growth of the indicated clones was monitored in the stem at the site of infection, 5 cm above the site of infection, and 10 cm above the site of infection at 7 and 14 dpi. Data are shown in a box plot, representing at least 19 replicates pooled from two experiments. (**D**, **E**) Bacteria were isolated and quantified from pooled terminal leaflet samples from the second, third, and fourth true leaves of each plant at 7 and 14 dpi. Data represent at least 30 replicates pooled from four (7 dpi) or three (14 dpi) experimental repeats. (**D**) Stacked bar graphs represent the presence/absence of bacteria in the sampled leaflets at 7 and 14 dpi. (**E**) Bar graphs show the averages and standard errors of bacterial populations in the leaves at 7 and 14 dpi. (**F**) Cm WT and CmΩ1284 were spotted (OD600 = 1) on LB agar supplemented with 5% sucrose and incubated for four days. Plates were photographed at 5 dpi (lower panel), and EPS production was quantified in the resulting clones (upper panel) using phenol– sulfuric acid hydrolysis assays followed by colorimetric quantification, standardized to bacterial count. (**G**) Cm WT and CmΩ1284 were grown on LB medium, standardized to OD600 = 0.5, and incubated in 24-well plates without shaking for 5 days. Supernatants were removed, wells were washed three times with distilled water, and attached bacteria were stained with crystal violet to quantify surface attachment (lower panel), followed by colorimetric quantification (upper panel). (**H**) Cm WT and CmΩ1284 were spotted (OD600 = 1) onto M9 medium supplemented with 0.1% carboxymethyl cellulose (CMC), and incubated for four days. Plates were stained with 0.1% Congo red, photographed (lower panel), and halo diameters were measured with a ruler to assess CMC degradation (upper panel). (**F**, **G**, **H**) Graphs represent means ± SE of at least ten biological replicates pooled from two experiments. (**B**–**H**) Asterisks (*) indicate that data were significantly different compared to Cm WT (U-test for C, E, F, G, and H; chi-squared test for B and D; p-value < 0.05).

These results support CMM_1284 as a regulator of the vascular lifestyle and its associated traits.

### Media-adapted clones do not demonstrate directional phenotypic selection

To confirm that tissue adaptation was due to host selective pressure rather than culture conditions, a control population of ten clones was generated by subculturing on LB agar for 15 passages. These clones (LB1–LB10) were analyzed for EPS production, cellulase activity, and virulence. Unlike tissue-adapted clones, which showed consistent shifts, media-adapted clones displayed scattered phenotypes: reduced EPS and cellulase activity in LB1 and LB3, complete cellulase loss in LB8, and altered virulence in LB9 and LB10 (Fig. S7). Thus, prolonged media culturing may cause mutations but not consistent, directional adaptation, likely reflecting clonal drift.

## Discussion

Bacterial pathogens switch strategies within their host to complete the infection cycle, involving changes in behavior, target tissues, and movement. To adapt to different infection phases, they must overcome diverse challenges and exhibit phenotypic plasticity. We investigated this plasticity by subjecting Cm to tissue-specific experimental evolution, differentiating adaptation to vascular versus apoplastic lifestyles. Ten parallel clones in each condition diverged phenotypically: vascular-adapted clones showed reduced virulence, lower cellulase activity, decreased EPS production, and enhanced surface attachment, whereas apoplast-adapted clones retained virulence, increased EPS production, and reduced surface attachment. These results indicate that xylem and apoplast exert distinct selective pressures, driving directional selection of traits. During disease, Cm exits xylem vessels late in infection, associated with symptom onset. Xylem residence may represent a dormant stage enabling systemic spread without symptoms, while apoplastic colonization likely drives vascular collapse and tissue maceration, in a similar manner to previous reports on *Erwinia amylovora* [8].

The molecular signals governing this shift are not well defined. In other xylem pathogens such as *E. amylovora* and *Ralstonia* spp., shifts are linked to population sensing and host/environmental cues like metabolites, physiology, and temperature [34,35]. It is possible that during vascular adaptation in this study, we selected clones that linger longer in the dormant stage, resulting in attenuated virulence. Traits consistently observed in tissue-adapted clones suggest strong selection on EPS production and surface attachment, which appear negatively correlated in Cm. In the apoplast, higher EPS may protect against antimicrobial responses [36–38], while vascular adaptation favors surface attachment and biofilm formation for xylem colonization [10]. Repeated vascular exposure should thus select for strong attachment and dense biofilms, redundant in the apoplast.

Notably, clones with higher surface attachment produced less EPS. Although EPS usually supports biofilms [31], composition and viscosity often matter more than quantity [32]. For instance, in *Xylella fastidiosa*, mutants lacking EPS-degrading enzymes overproduce EPS, leading to reduced surface attachment and impaired biofilm formation—similar to what we observed in apoplast-adapted clones [39]. EPS overproduction may inhibit cell-to-cell or surface attachment by masking cell wall-anchored proteins [40]. Notably, in many Gram-positive Firmicutes, surface attachment and biofilm formation are mediated by cell wall-bound proteins rather than EPS [40–42]. This suggests that EPS may not contribute significantly to biofilm formation in at least some Gram-positive species, contrasting with widely accepted Gram-negative-based models. This likely underlies attenuated virulence: vascular-adapted clones showed reduced stem-to-leaf migration, possibly due to impaired biofilm disassembly, and decreased vessel blockage, as EPS strongly contributes to wilting. Dynamic EPS modulation thus appears central to Cm colonization: low EPS supports local xylem establishment, while increased EPS promotes dispersal and systemic spread. These transitions are probably regulated by physiological and environmental cues. Another trait in vascular-adapted clones was reduced cellulase and amylase activity. Cellulase, linked to virulence through the secreted glucanase CelA, is essential for symptom development [20,33]. Cell wall–degrading enzymes can also affect EPS and biofilm disassembly [43,44], though not for CelA: *celA* mutants lack EPS phenotypes, overexpression does not restore vascular clone EPS traits, and natural isolates show no correlation between cellulase, morphology, or EPS. Therefore, CelA more likely modulates plant cell walls and xylem exit, promoting spread as in *Pectobacterium* [45]. Overexpressing *celA* only partially restored virulence in vascular clones, suggesting phenotypes depend more on EPS and surface attachment. Why cellulase was negatively selected remains unclear and warrants further study.

Despite major phenotypic changes, adapted clones accumulated few SNPs/INDELs, with no large deletions or plasmid loss. Mutation frequencies align with previous studies [46], though stress may elevate rates via reactive oxygen species [47]. Importantly, neither *celA*, its promoter, pCM1, nor EPS-associated clusters were mutated, implying upstream regulatory changes. Indeed, six of ten vascular clones fixed mutations the in transcriptional regulator *CMM_1284*, causing amino acid substitutions in its DNA-binding domain. A marker exchange mutant of *CMM_1284* recapitulated vascular-clone phenotypes: enhanced attachment, reduced EPS and cellulase activity, and attenuated virulence, suggesting a role in vascular–apoplastic shifts. However, some clones lacked *CMM_1284* changes yet showed similar phenotypes, and two vascular clones had no coding mutations at all, indicating regulatory, structural, or epigenetic mechanisms undetected by Illumina sequencing.

In summary, experimental evolution identified tissue-specific traits in Cm. Biofilm formation and surface attachment were critical for vascular colonization but limited migration and virulence, suggesting that biofilm disassembly is essential for transitioning from a latent vascular phase to a virulent apoplastic phase.

## Materials and methods

### Bacterial Strains and Plant Material

Cm and *E. coli* strains developed and used in this study are listed in Table S2. Cm strains were grown in Luria-Bertani (LB) medium at 28°C, supplemented with 100 µg/ml trimethoprim [48]. *E. coli* strains were grown in LB medium at 37°C. When required, media were supplemented with 25 µg/ml gentamicin, 75 µg/ml neomycin, 50 µg/ml kanamycin, or 100 µg/ml ampicillin. The plant cultivars used in this study were tomato (*Solanum lycopersicum* cv. Moneymaker), eggplant (*Solanum melongena* cv. Black Queen), and *Nicotiana sylvestris*. Plants were grown in a temperature-controlled glasshouse at 25°C under natural light conditions.

### Experimental Evolution Procedure

Vascular, apoplastic, and media-adapted populations were generated by repeated passaging of Cm NCPPB382 in tomato stems, leaves, and LB medium. A full description is provided in Supplementary File 1. Briefly, for vascular passages, bacteria were toothpick-inoculated into stems of 4–6 leaf plants and re-isolated ∼6 cm above the inoculation site after 14–21 days. For apoplastic passages, diluted suspensions (10□ CFU/ml) were syringe-infiltrated into mature leaves and recovered from pooled foci after 7 days. Ten parallel lines per tissue type were passaged for 15 cycles, with populations stored in glycerol at –80 °C. Representative clones from passages 5, 10, and 15 were used for phenotyping and sequencing.

### Plant inoculations, disease severity assessments, apoplastic virulence, stem-to-leaf migration and quantification of stem/leaf bacterial populations

The virulence assays were carried out using wound inoculation method similar to that described by [49], with some minor adjustments. A full description is provided in Supplementary File 1.

Virulence assays were conducted using toothpick-mediated stem wound inoculation and needleless syringe infiltration of tomato leaves. Wilt symptoms in stems were scored at 14 or 21 dpi on a 0–3 scale based on the percentage of affected leaves. Leaf infiltration assays were evaluated at 10 dpi for chlorosis and necrosis on a 0–3 scale, and representative leaves were photographed. Bacterial populations were quantified from stem segments or leaf disks by plating serial dilutions, standardized to tissue weight or surface area. Hypersensitive response assays were performed in eggplant and *Nicotiana sylvestris* by syringe infiltration, with symptoms recorded at 36 h.

### Cloning and Bacterial Manipulation

All plasmids and oligonucleotides produced and used in this study are listed in Tables S3 and S4, respectively. Detailed information regarding cloning techniques, and plasmids construction is provided in Supplementary File 1. The plasmid pMA-RQ:Cmp [18] served as the backbone for constructing marker exchange, overexpression, and reporter plasmids. A marker exchange construct (pCMAT:1284) targeting *CMM_1284* was generated to obtain deletion mutants, which were confirmed by PCR. For overexpression, the backbone was modified with a promoter and a genomic integration site, producing derivatives such as pCMIARG, including a CelA-3×HA fusion confirmed by western blot. Reporter plasmids carrying *celA* and *gyrB* promoters fused to the *uidA* (GUS) gene were built in the Cm–*E. coli* shuttle vector pHN216 [50] and used for promoter activity assays. All constructs were introduced into Cm by electroporation, as described in [49].

### EPS quantification, surface attachment, and plate halo assays

Detailed descriptions of experimental procedures used for in vitro characterization is provided in Supplementary File 1. Briefly, EPS quantification was conducted on Cm bacteria grown in 5% supplanted LB plates, using the phenol-sulfuric acid estimation method [51]. Surface attachment/biofilm assays were performed using 24 well plates supplemented with Cm cultures previously grown on LB broth. Attachment was assessed and quantified after static incubation using crystal violet staining method as previously described [52] with modifications. Exoenzyme activity of Cm clones was tested by plate halo assays on media containing specific substrates for cellulase, amylase, xylanase, polygalacturonase, protease, or lipase, with halos visualized by staining and quantified by diameter measurements [53]. Siderophore production was assessed using the CAS agar assay [54] under iron-limiting conditions, with activity visualized as yellow/orange halos around colonies.

### *In Situ* Localization of *Clavibacter michiganensis* During Infection Using Confocal Laser Scanning Microscopy

Cm clones NCPPB382, S2, and S4 were transformed with the *E. coli*–*Clavibacter* shuttle vector pK2-22 [24], which expresses EGFP under the control of the p*CMP1* promoter. *In situ* localization of bacteria was monitored in infected plants using an Olympus IX81 confocal laser scanning microscope equipped with a GFP filter set. To visualize bacterial localization in stem tissues, ∼0.1□mm-thick horizontal and vertical cross-sections were collected approximately 3□cm above the inoculation site. For leaf tissues, ∼0.5□cm² sections were excised from areas exhibiting healthy, wilted, or necrotic phenotypes and directly mounted on microscope slides for imaging.

### Promoter Activity Assays and Gene Expression Analysis

For promoter activity and gene expression assays, Cm WT, S2, S4, and S9 strains, either alone or carrying pHN:p*celA*:*GUS* or pHN:*pgyrB*:*GUS*, were grown overnight in LB, washed twice with distilled water, resuspended in M9 medium to induce virulence-associated gene expression [55], and incubated at 28 °C with rotation for 24 h.

For promoter activity, OD_600_ was measured, and 1.5 ml of each culture was lysed with 1 mg/ml lysozyme followed by sonication (SONIC-150W, MRC Labs). Lysates were centrifuged, and 80 µl of supernatant was mixed with 10 µl 10× Tris-based saline (pH 7.4) and 10 µl 10 mM PNPG. Reactions were incubated at 37 °C and monitored for yellow pigmentation, then stopped with 50 µl 1 M sodium bicarbonate. Absorbance at 405 nm was measured, and activity was calculated as OD_405_ / (time × OD_600_) and standardized to WT activity.

For gene expression, total RNA was extracted from culture supernatants using the Hybrid-R Kit (GeneAll), reverse transcribed with the UltraScript™ cDNA Synthesis Kit (PCR Biosystems), and amplified using Fast SYBR qPCR Master Mix (Biogate) on a QuantStudio 3 system (Applied Biosystems) with gene-specific primers (Table S3). Expression was normalized to *gyrA* and calculated with the comparative Ct method [56].

### Genome Sequencing, Assembly, and Comparative Genomic Analysis

Three independent Cm NCPPB382 (WT) clones and tissue-adapted clones collected after 5, 10, and 15 passages were subjected to whole-genome sequencing. DNA was extracted from 10 ml overnight LB cultures using the Wizard Genomic DNA Purification Kit (Promega). Libraries were sequenced on an Illumina NextSeq2000 (150 bp paired-end) at the Applied Microbiology Services Laboratory (Ohio State University). Reads were cleaned with Trimmomatic [57] and assembled using Unicycler v0.5.0 and SPAdes v3.15.5 [58,59]; contigs <200 bp were removed. Genome completeness was assessed with BUSCO v5 (micrococcales_odb10) [60]. Genomes were deposited in the NCBI BioProject database under the ID PRJNA1333927. Genomic alterations were identified with Snippy (https://github.com/tseemann/snippy) and confirmed by BWA [61] against the NCPPB382 reference (GCA_000063485). Passage 10 of clone S6 and passage 5 of clone S10 were excluded due to poor sequencing quality. All INDELs and SNPs are listed in Table S1, excluding mutations found in any of the three resequenced WT clones. Ten representative ORF mutations were confirmed by Sanger sequencing (Table S1, *). The corresponding regions were PCR-amplified from Cm WT and adapted clones using specific primers (Table S4), sequenced (Hylabs Laboratories), and compared with assembled genomes.

## Acknowledgments

This work was supported by the Israel Binational Science Foundation (BSF, grant no. 2021190 and 2023281, for D.T. and J.M.J).

## Notes

### Competing Interest Statement

The authors have declared no competing interest.

