## Supplementary material for "Tissue-Specific Experimental Evolution Reveals Adaptive Trade-Offs in the Plant Vascular Pathogen *Clavibacter michiganensis*": S: S1.docx

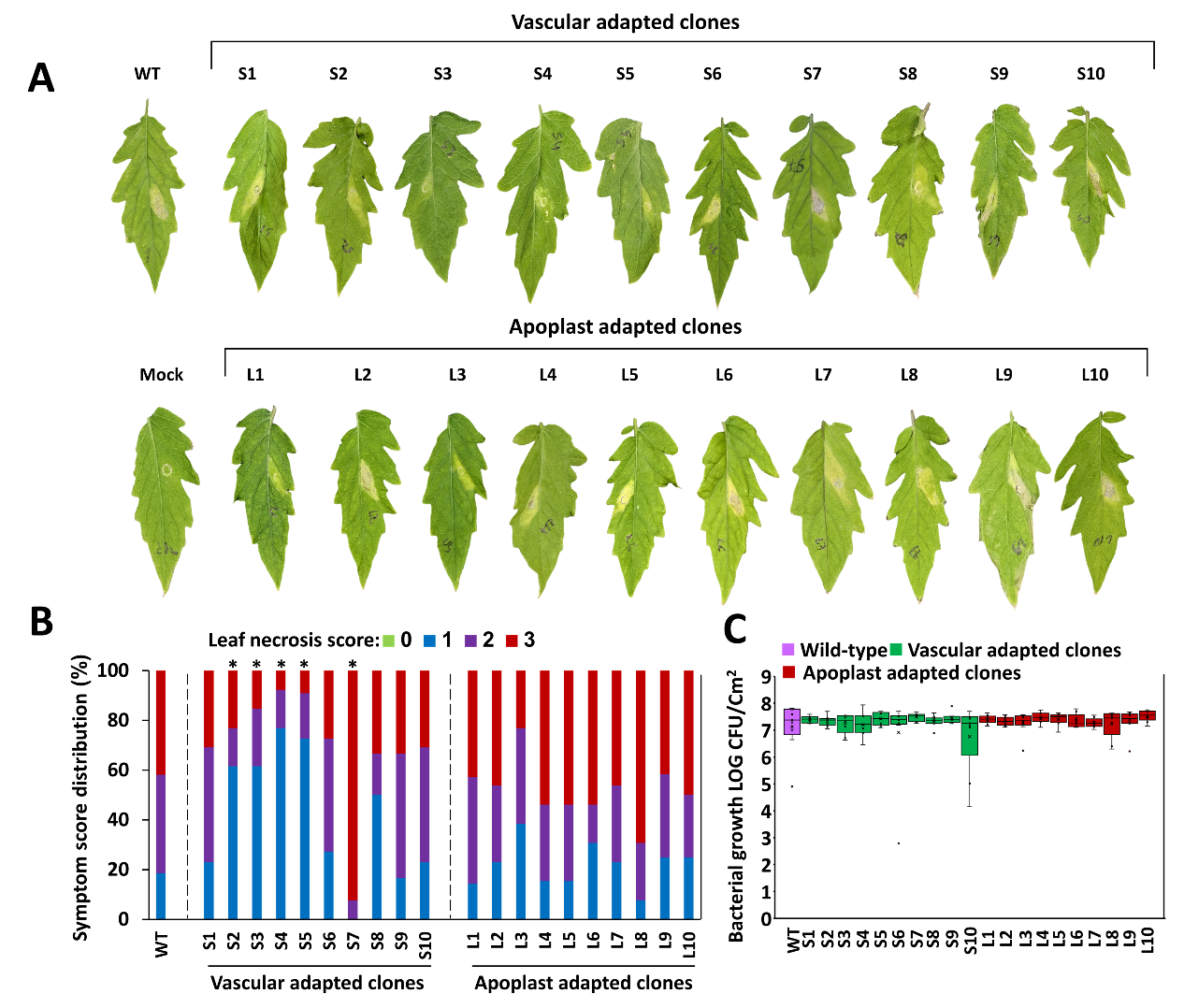

**Fig. S1. Vascular adaptations affect apoplastic virulence.** Six-leaf stage tomato leaves were inoculated with the indicated vascular adapted clones (S1-S10), apoplast adapted clones (L1-L10) and Cm382 (WT) through infiltration of bacterial cultures (10^4^ CFU/ml) using a needless syringe. (**A)** Representative leaves (out of at least 11 repeats) were photographed at 10 days post-infiltration (dpi). **(B)** Necrotic symptoms were scored at 10 dpi according to the following scale: 0 – no symptoms, 1 = chlorosis alone, 2= necrosis of 1-50% of the infected area, 3= necrosis of 51-100% of the infected area. The graph depicts the distribution of at least 11 repeats for each clone pooled from two experiments. "*" represent significant difference (chi-squared test, p-value < 0.05) compared to WT. (**C)** Bacterial growth in the infiltration sites at 10 dpi. Data represents 10 repeats for each clone pooled from two experiments. No significant differences (U-test) were observed between each of the clones to the WT.
