## Supplementary material for "Tissue-Specific Experimental Evolution Reveals Adaptive Trade-Offs in the Plant Vascular Pathogen *Clavibacter michiganensis*": S: S2.docx

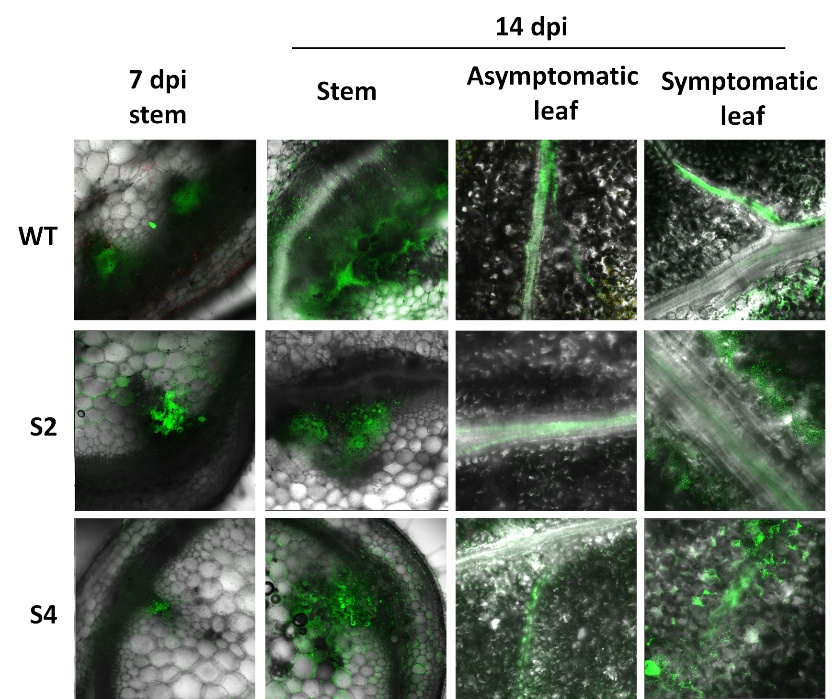

**Fig. S2. *In situ* localization of vascular-adapted Cm during infection**. Stem areas between the cotyledons of four-leaf-stage tomato plants were inoculated by wounding with toothpicks soaked in GFP-labeled bacterial suspensions (10^7^ CFU/ml) of Cm WT and the vascular-adapted clones S2 and S4. Bacteria were visualized in horizontal stem cross-sections, as well as in asymptomatic and symptomatic leaves, at 7 and 14 days post-inoculation. The data shown are representative of at least ten biological replicates conducted across three independent experiments.
