## Supplementary material for "Tissue-Specific Experimental Evolution Reveals Adaptive Trade-Offs in the Plant Vascular Pathogen *Clavibacter michiganensis*": S: S3.docx

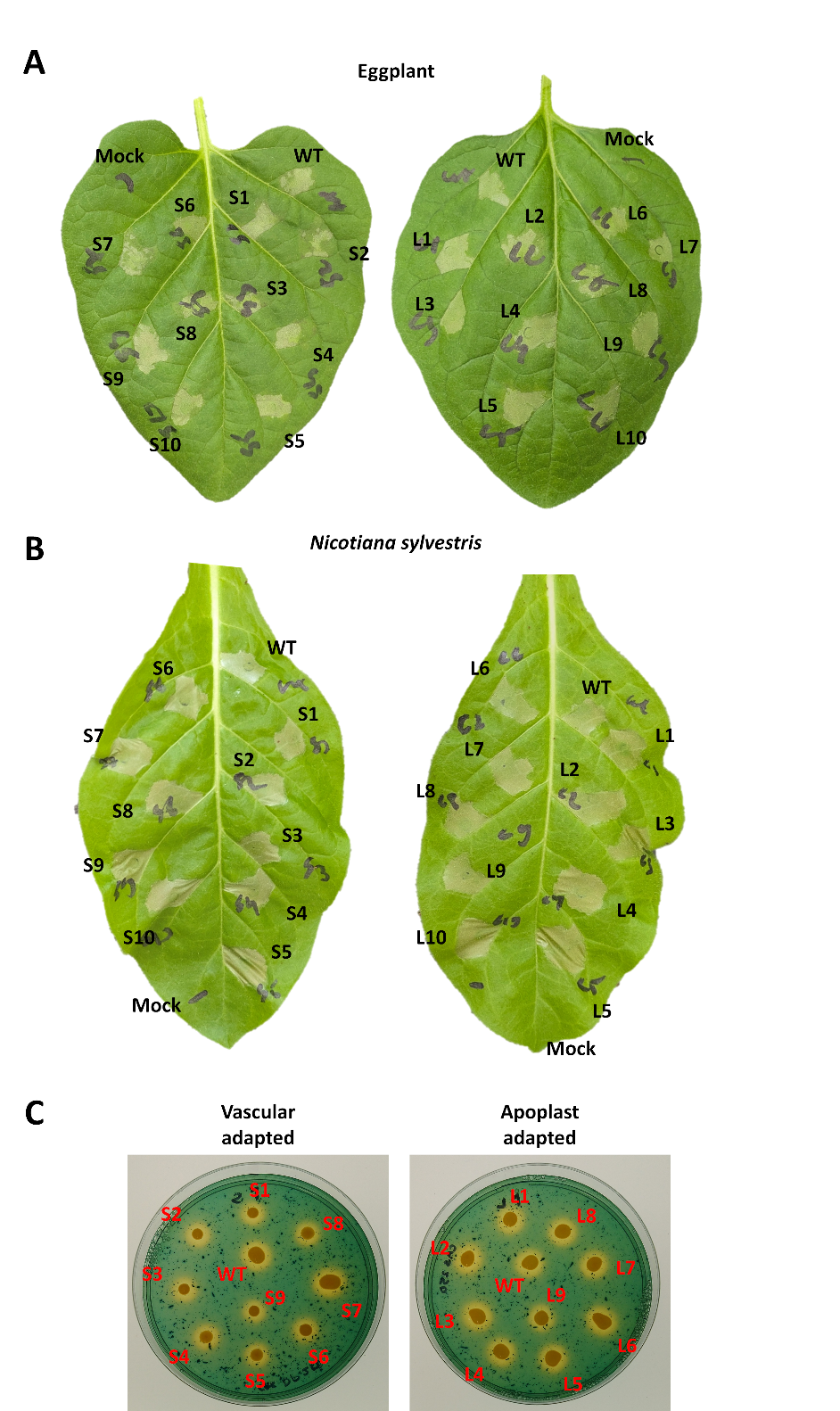

**Fig. S3. Vascular and apoplastic adaptations do not affect hypersensitive response induction and siderophore production**. (**A,** **B**) Leaves of six‑leaf‑stage eggplant (**A**) or *Nicotiana sylvestris* (**B**) plants were infiltrated with Cm WT and the indicated vascular and apoplast adapted clones using a needleless syringe at a concentration of 5 × 10^7^ CFU/ml. Representative leaves were photographed 36 h post‑infiltration. (**C**) Cm WT and the indicated vascular and apoplast adapted clones were spotted (OD600=1) on LB agar supplemented with CAS indicator dye and 50 µM 2,2'-dipyridyl and incubated for ten days and photographed. Experiments were repeated twice, with at least five independent replicates per experiment, yielding consistent results.
