## Supplementary material for "Tissue-Specific Experimental Evolution Reveals Adaptive Trade-Offs in the Plant Vascular Pathogen *Clavibacter michiganensis*": S: S4.docx

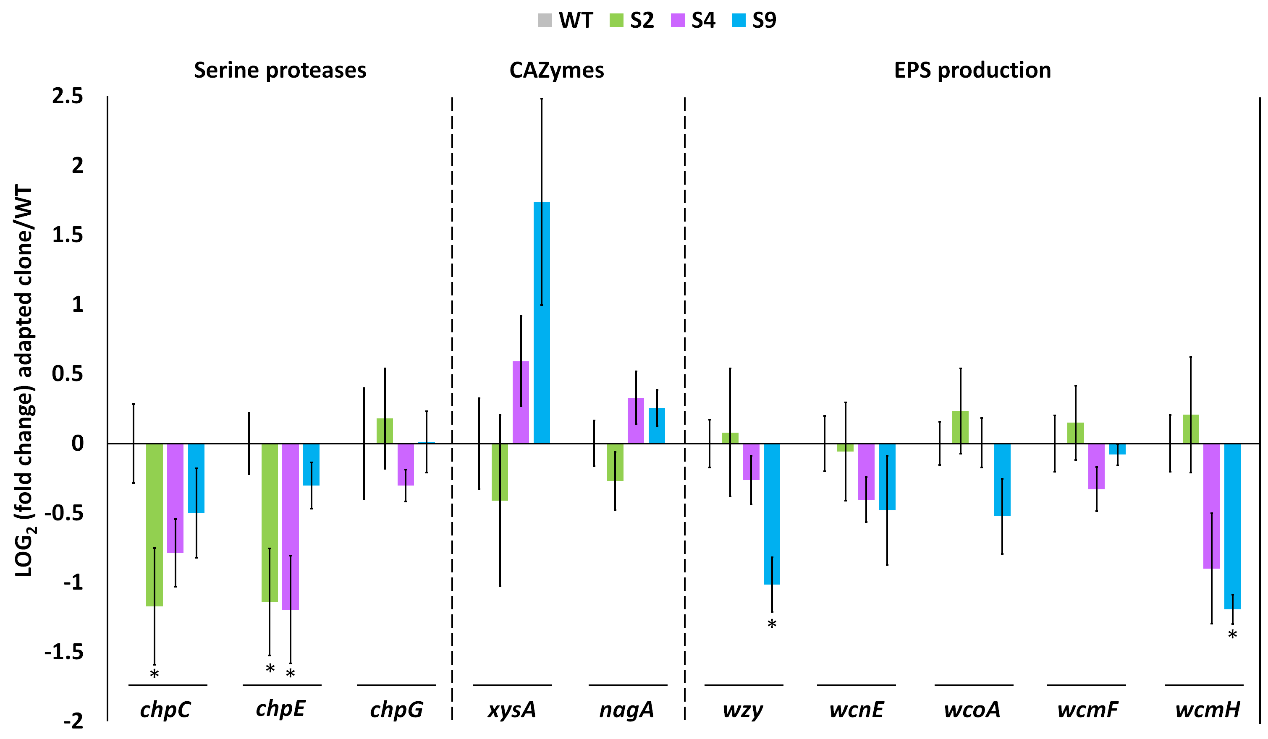

**Fig. S4.** **Transcriptional expression of virulence-associated genes in vascular-adapted clones**. mRNA transcript abundance was quantified by RT‑qPCR for the putative serine protease genes chpC (CMM_0052), chpE (CMM_0039), and chpG (CMM_0059), the putative CAZyme genes xysA (CMM_1673) and nagA (CMM_0049), and genes predicted to be associated with EPS production, including wzy (CMM_0715), wcnE (CMM_0718), wcoA (CMM_0819), wcmF (CMM_1597), and wcmH (CMM_1601). Cultures were incubated for 24 h in sucrose‑supplemented M9 medium. gyrA (CMM_0007) was used for normalization. Graph depict the mean ± SE of relative transcript abundance compared to Cm WT, based on six independent biological replicates pooled from two independent experiments. Asterisks (*) indicate statistically significant differences versus WT (U-test, p-value < 0.05).
