## Supplementary material for "Tissue-Specific Experimental Evolution Reveals Adaptive Trade-Offs in the Plant Vascular Pathogen *Clavibacter michiganensis*": S: S5.docx

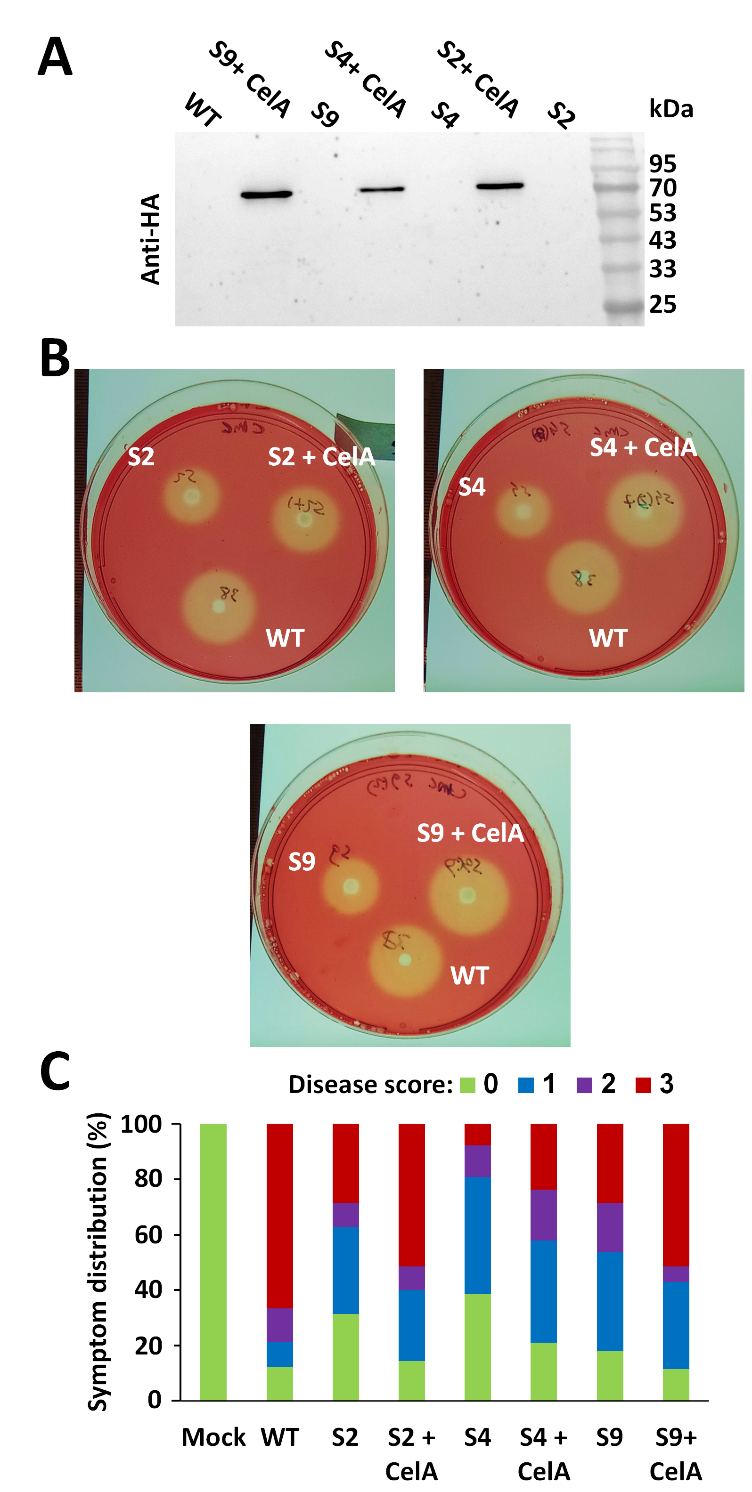

**Fig. S5. Effect of *celA* overexpression in vascular-adapted clones on exocellulase activity and vascular virulence**. An integrative plasmid containing *celA* fused to an HA tag under the control of the *MegroEL* promoter was introduced into vascular-adapted clones S2, S4, and S9. Introduction of this construct is indicated as “+ CelA”. (**A**) CelA-HA protein accumulation in the supernatants of the indicated overnight-grown Cm cultures was detected by western blot using an HA-specific antibody. (**B**) The indicated Cm cultures (OD600 = 1) were spotted onto M9 medium supplemented with 0.1% carboxymethyl cellulose (CMC) and incubated for four days. Plates were stained with 0.1% Congo red and photographed. (**C**) Four-leaf-stage tomato plants were wound-inoculated with the indicated Cm clones at the stem area between the cotyledons. Wilt symptoms were scored as the percentage of wilted leaves using the following scale: 0 = no wilting, 1 = 1–25%, 2 = 25–50%, 3 = 51–100%. The stacked graph shows the distribution of wilt scores for 24 replicates per clone, pooled from three independent experiments at 21 dpi.
