## Supplementary material for "Tissue-Specific Experimental Evolution Reveals Adaptive Trade-Offs in the Plant Vascular Pathogen *Clavibacter michiganensis*": S: S6.docx

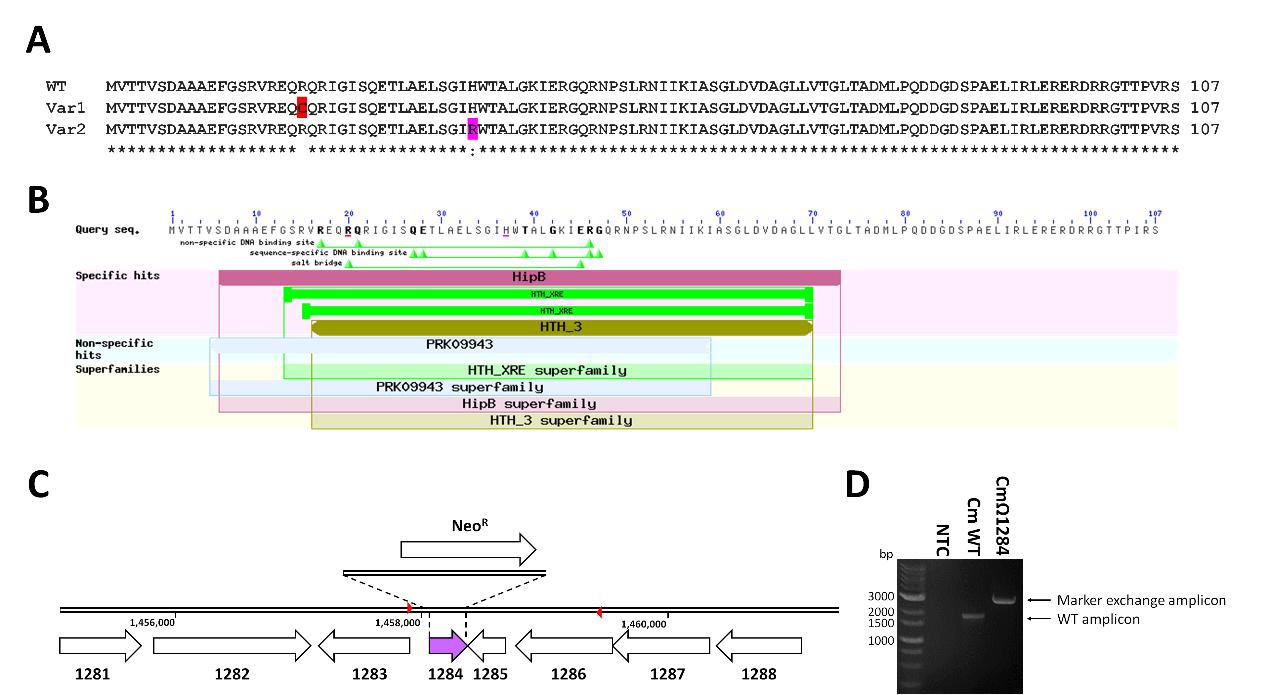

**Fig. S6. Sequence analysis of CMM_1284 and confirmation of the CMM_1284 marker-exchange mutant.** (**A**) Protein sequence alignment of Cm NCPPB382 CMM_1284 and its modified variants Var1 (found in S1, S2, and S6; amino acid substitution is marked in red) and Var2 (found in S3, S4, and S9; amino acid substitution is marked in magenta) in vascular-adapted clones. The alignment was performed using Clustal Omega (<https://www.ebi.ac.uk/jdispatcher/msa/clustalo>). (**B**) Domain prediction of CMM_1284 using the NCBI conserved domain feature (<https://www.ncbi.nlm.nih.gov/Structure/cdd/wrpsb.cgi>). Amino acid substitutions present in CMM_1284 Var1 and Var2 are marked in red and magenta, respectively. (**C**) Physical map of the genomic region surrounding *CMM_1284* (1,455,000–1,461,500 bp) on the NCPPB382 bacterial chromosome (NC_009480.1). The *CMM_1284* gene is marked in purple. The region replaced by the neomycin resistance gene is indicated by a dotted line. The primers used for confirmation of the gene replacement shown in (**D**) are indicated by red arrows. (**D**) PCR confirmation of *CMM_1284* gene replacement in the marker-exchange mutant CmΩ1284.
