## Supplementary material for "Tissue-Specific Experimental Evolution Reveals Adaptive Trade-Offs in the Plant Vascular Pathogen *Clavibacter michiganensis*": S: S7.docx

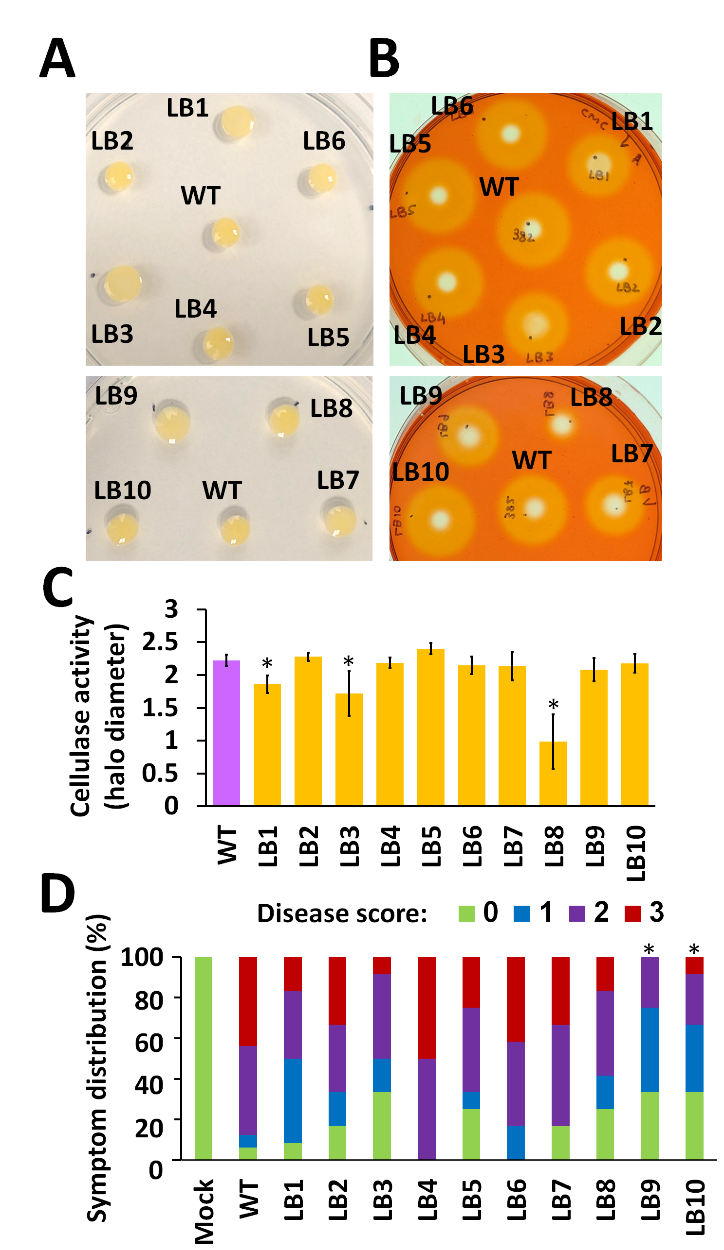

**Fig. S7. Characterization of LB culture-adapted Cm clones.**  (**A**) Cm WT and the LB media adapted clones (LB1-LB10) were spotted (OD600 = 1) on LB agar supplemented with 5% sucrose and incubated for four days. Plates were photographed at 5 dpi. (**B**, **C**) Cm WT and the indicated clones were spotted (OD600 = 1) onto M9 medium supplemented with carboxymethyl cellulose (CMC) ,and 0.01% sucrose, then incubated for four days. Plates were stained with 0.1% Congo red, photographed (**B**), and halo diameters were measured with a ruler to assess CMC degradation (**C**). Asterisks (*) indicate that data were significantly different compared to Cm WT (U-test, p-value < 0.05). (**D**) Four-leaf-stage tomato plants were wound-inoculated with the indicated Cm clones at the stem area between the cotyledons. Wilt symptoms were scored as the percentage of wilted leaves using the following scale: 0 = no wilting, 1 = 1–25%, 2 = 25–50%, 3 = 51–100%. The stacked graph shows the distribution of wilt scores per clone, at 14 dpi. Asterisks (*) indicate that data were significantly different compared to Cm WT (chi-squared test, p-value < 0.05). All data represent at least seven (A, B, C) or 12 (D) biological repeats pooled from two independent experiments.
