## Supplementary material for "Tissue-Specific Experimental Evolution Reveals Adaptive Trade-Offs in the Plant Vascular Pathogen *Clavibacter michiganensis*": S: Supplementary file 1.docx

**Supplementary file 1. Extended Materials and Methods**

**Experimental Evolution Procedure**
Vascular and apoplastic populations were generated through repeated passaging of Cm in tomato stems and leaves, using Cm NCPPB382 as the parent strain. For each **vascular passage**, bacteria isolated from the plant in the previous round were inoculated into a naïve four-to-six leaf stage tomato plant via a single stem puncture (between the cotyledons) using a toothpick previously incubated in a bacterial suspension (10⁷ CFU/ml; “wound inoculation” method). Plants were then transferred to a growth room, and 14–21 days later, bacteria were re-isolated from 3 mm stem sections located ~6 cm above the inoculation site. For each **apoplastic passage**, bacterial suspensions from the previous round were diluted to 10^4^ CFU/ml and infiltrated into the three most recently emerged mature leaves using a needleless syringe. Each leaf received 4–8 infiltration sites (~1 cm² per site). Plants were incubated in a growth room for 7 days, after which bacteria were isolated from pooled samples of all infection foci. For each vascular and apoplastic adaptation cycle, bacteria were isolated by homogenization, serial dilution, and plating on LB agar supplemented with 50 µg/ml cycloheximide and 100 µg/ml trimethoprim. Approximately 50–200 colonies from each sample were pooled and used for the subsequent passage. The experiment was carried out using ten parallel bacterial clone lines for each tissue type (vascular and apoplastic), with a total of 15 passages. Each passage was performed in a separate plant (20 plants per passage), and bacterial colonization was quantified at each time point (CFU per gram of stem tissue or per cm² of leaf tissue). Adapted bacterial populations from each passage were stored in 25% glycerol at –80 °C. For downstream phenotyping and genomic sequencing, bacteria from all parallel lines at passages 5, 10, and 15 were streaked on LB agar, and a single representative colony from each clone was selected. Clones from passage 15 were used for phenotyping, and those from passages 5, 10, and 15 were used for sequencing.

Culture-media adaptation was conducted by pooling bacteria grown in culture media from the previous passage in 1 ml of distilled water, diluting the suspension to OD600 = 0.001, and spreading it on LB agar supplemented with trimethoprim. Plates were incubated for four days and then pooled for the next passage. Media passaging was performed on ten parallel clones for 15 cycles.

**Plant inoculations, disease severity assessments, apoplastic virulence, stem-to-leaf migration and quantification of stem/leaf bacterial populations**

The virulence assays were carried out using wound inoculation method similar to that described by [1], with some minor adjustments. Stem "wound inoculations" were conducted by a single punctures of the stem areas between the cotyledons of four-leaf tomatoes with pre-soaked (10 minute) Cm-contaminated (10^7^ CFU/ml) toothpicks. After inoculations, plants were kept at 25˚C in a glasshouse under natural light conditions and wilt symptoms were determined and scored at 14 or 21 dpi. Wilting symptoms were scored in each plant as the percentage of leaves demonstrating wilting and/or necrotic blotch symptoms according to the following scale: 0 = no wilt or leaf blotch, 1 = 1%–25%, 2 = 26%–50%, 3 = 51%–100%.

To determine the effect of tissue adaptation on apoplastic virulence, six-leaf stage tomato leaves were inoculated through infiltration of bacterial cultures (10^4^ CFU/ml) using a needleless syringe. Representative leaves were photographed at 10 days post-infiltration (dpi). Necrotic symptoms were scored at 10 dpi according to the following scale: 0 – no symptoms, 1 = chlorosis alone, 2= necrosis of 1-50% of the infiltrated area, 3= necrosis of 51-100% of the infiltrated area.

Bacterial quantification in stem tissues was performed by collecting 1-mm stem segments and determining their fresh weight. For leaf tissues, 0.5-cm diameter leaf disks were collected either from the infection site (for infiltration assays) or from the central midveins of the terminal leaflets of the second, third, and fourth true leaves (for leaf migration assays). Samples were homogenized, and bacterial populations were quantified by plating 10 μl of 10-fold serial dilutions and counting the resulting colonies. Values were standardized based on tissue weight (for stems) or surface area (for leaves).

For hypersensitive response (HR) visualization, leaves of six‑leaf‑stage eggplant or *Nicotiana sylvestris* plants were infiltrated with the bacterial cultures (5 × 10^7^ CFU/ml) using a needleless syringe. Representative leaves were photographed 36 h post‑infiltration.

**Cloning and Bacterial Manipulation**

All plasmids and oligonucleotides produced and used in this study are listed in Tables S3 and S4, respectively. All constructs were introduced into Cm by electroporation, as described in [1]. The plasmid pMA-RQ:Cmp (originally carrying the p*CMP1* promoter–MCS–3×HA cassette [2]) was used as a backbone for constructing marker exchange and overexpression plasmids. Inserts were generated by amplifying the desired fragments using Phanta Flash Super-Fidelity DNA Polymerase (Vazyme) from templates including the plasmids pHN216 [3] and pSelACT-KO [4](carrying ***nptII*** and ***aac(3)-IV***, respectively), and genomic DNA from Cm NCPPB382, Microbacterium esteraromaticum, and E. coli. Fragments were cloned into the desired constructs by either restriction-based cloning or using the LigON Kit (EURX).

To produce a marker exchange plasmid, the gentamicin resistance gene ***aac(3)-IV*** was cloned upstream of the p*CMP1* promoter in pMA-RQ:Cmp to generate pCMAT. A marker exchange plasmid targeting ***CMM_1284*** was constructed by introducing a cassette composed of a 498 bp 5′ flanking region (positions 1457574–1458071 in AM711867), the ***nptII*** gene (neomycin resistance), and an 852 bp 3′ flanking region (positions 1458369–1459232 in AM711867) into pCMAT, producing pCMAT:1284.

To generate the ***CMM_1284*** marker exchange mutant, pCMAT:1284 was introduced into Cm NCPPB382. Transformed bacteria were plated on LB medium supplemented with neomycin, and resulting colonies were double-patched onto LB plates containing either neomycin alone or neomycin plus gentamicin to identify double-crossover events. Clones that retained neomycin resistance but failed to grow on gentamicin were verified by PCR using primers targeting the flanking regions of ***CMM_1284*** (Table S4, Fig. S5C and S5D).

To construct the integrative overexpression plasmid, pCMAT was further modified by replacing the p*CMP1* promoter with the –228 to +3 bp 5′ putative promoter fragment of the ***groEL*** gene (pMegroEL, positions 289679–289911 in CP043732; locus tag FVO59_01375 from M. esteraromaticum). A homologous recombination site in the 455 bp 3′ untranslated region of the ***chpB*** (CMM_PS_10) pseudogene (positions 89863–90317 in AM711867) was added downstream of the 3×HA tag, resulting in plasmid pCMIARG. For ***celA*** overexpression, the ***celA*** gene (pCM1_0020) was cloned in-frame with the 3×HA tag downstream of p*MegroEL* and introduced into Cm clones NCPPB382, S2, S4, and S9. Protein accumulation of CelA was confirmed by western blot in overnight grown culture using anti-HA-tag (F-7) mouse monoclonal antibody (Santa Cruz Biotechnology) as described by [1,5] according to the manufacturer's instructions.

To generate GUS reporter plasmids for promoter activity assays, the putative promoters of ***celA*** (pCM1_0020, positions 16938–17653 in AM711865) and ***gyrB*** (CMM_0006, positions 5324–5957 in AM711867) were fused to the E. coli ***uidA*** gene by overlap extension PCR. The resulting fragments were cloned into the E. coli–Cm shuttle vector pHN216 (OR234300, [3]) and introduced into Cm clones NCPPB382, S2, S4, and S9. Reporter-based GUS activity was confirmed by patching transformants and their parental clones in parallel onto LB plates supplemented with X-gluc. Blue pigmentation appeared within 4–7 days in all transformants, but not in any of the parental clones.

**Separation and Quantification of EPS**

EPS were isolated from the cell-free culture extract of Cm strains using phenol-sulfuric acid estimation method [6]. Overnight-grown Cm cultures were pelleted by centrifugation and washed twice with double-distilled water. Washed bacterial suspensions were adjusted to OD600 = 1, and 15 µl aliquots were spotted onto LB plates supplemented with 5% sucrose. Plates were incubated at 28° C under static conditions for 6 days. Bacterial colonies were then scraped and resuspended in 1 ml of double-distilled water. From each suspension, 100 µl was used for serial dilution plating to determine CFU. The remaining suspension was centrifuged at 7,969 × g for 10 min to separate EPS from the cells. Ice-cold acetone (two volumes) was added to the supernatant, and the mixture was incubated overnight at 4°C. EPS was precipitated by centrifugation at 7,969 × g for 20 min, then dried at 37°C. EPS quantification was performed by measuring total carbohydrate content using the phenol–sulfuric acid colorimetric method, with D-glucose as the standard. EPS pellets were dissolved in 1 ml of Milli-Q water, and 400 µl of the solution was transferred to a test tube. Then, 200 µl of 5% (v/v) water-saturated phenol was added, followed by 1 ml of 44% concentrated H₂SO₄ added from the top with proper mixing. The mixtures were incubated at room temperature for 10 min, followed by incubation at 30°C for 20 min to develop an orange chromophore. Absorbance was measured at 490 nm, and carbohydrate content was calculated using a glucose standard curve. EPS values were normalized to CFU counts. The calibration curve was generated using a D-glucose stock solution (10 mg/ml) diluted to final concentrations ranging from 0 to 0.0275 mg/ml in 400 µl volumes.

**Surface Attachment Assays**
Surface attachment/biofilm assays were performed as previously described [7] with modifications. Cm strains were grown in LB broth supplemented with the appropriate antibiotics. Two milliliters of overnight-grown cultures were normalized to OD600=0.5 and transferred into sterile 24-well polystyrene culture plates. Plates were incubated statically at 28°C. After 10 days, the media was gently decanted, and the wells were washed twice with autoclaved water to remove loosely attached cells. Surface attachment was assessed using 1% crystal violet staining. Excess stain was removed by washing the wells with double-distilled water. Biofilm formation was quantified by dissolving the stained cells in 1 ml of 90% ethanol, and absorbance was measured at 570 nm.

**Exoenzyme Activity and Siderophore Production Assays**
Exoenzyme activity and siderophore production were assessed using plate halo assays. A 5 µl suspension of Cm clones, standardized to OD_600_= 1, was spotted onto LB or M9 agar plates supplemented with 0.1% sucrose, 0.25 g/L yeast extract, and 0.5 g/L Tryptone, containing appropriate substrates. Substrate degradation was photographed and quantified by measuring the diameter of the resulting halos.

Cellulase, amylase, xylanase, and polygalacturonase activities were assessed using modified M9 media containing 0.1% of carboxymethyl cellulose (CMC), starch, xylan, or polygalacturonate, respectively. Halo diameters were recorded at dpi for CMC and at 10 dpi for the other substrates. Plates were stained with 0.1% Congo red (for cellulase and xylanase), iodine (for amylase), or rhodamine B (for polygalacturonase). After staining, CMC and xylan plates were destained with 1 M NaCl to improve visualization. Protease and lipase activities were assayed on LB agar plates supplemented with 1% skim milk or 3% tributyrin, respectively. Substrate clearance was evaluated at 10 dpi. No protease or polygalacturonase activity was observed in any of the tested Cm clones.

Siderophore production by was assessed using the Chrome Azurol S (CAS) agar plate assay. CAS indicator dye (60.5 mg in 50 mL) was incorporated into LB agar supplemented with 50 µM 2,2'-dipyridyl to create iron-limiting conditions and allow visual detection of siderophore activity. Siderophore secretion, visualized as a yellow or orange halo around the bacterial colony, was monitored at 10 dpi.
