## Supplementary material for "Tissue-Specific Experimental Evolution Reveals Adaptive Trade-Offs in the Plant Vascular Pathogen *Clavibacter michiganensis*": S: Table 1.docx

**Table 1. Coding-sequence substitutions and short indels in vascular and apoplast-adapted clones**

| **Protein sequence changes** | **Putative annotation** | **Locus tag** | **Mutation** | **Clone** | **Type** |
| --- | --- | --- | --- | --- | --- |
| R20C | Transcriptional regulator, HipB/XRE-type | CMM_1284 | C1458114->T | S1 | **Vascular adaptation** |
| R20C | Transcriptional regulator, HipB/XRE-type | CMM_1284 | C1458114->T | S2 |  |
| H37R | Transcriptional regulator, HipB/XRE-type | CMM_1284 | A1458166->G | S3 |  |
| L158V | Hypothetical protein | CMM_2231 | C2524625->G |  |  |
| A328V | Two component sensor kinase | CMM_0996 | C1143378->T | S4 |  |
| H37R | Transcriptional regulator, HipB/XRE-type | CMM_1284 | A1458166->G |  |  |
| P68S | Metallo-beta-lactamase | CMM_1971 | A2227810->G |  |  |
| R20C | Transcriptional regulator, HipB/XRE-type | CMM_1284 | C1458114->T | S6 |  |
| A118->AAA | Metal-transporting ATPase | CMM_1037 | C1187404->CGCTGCT | S7 |  |
| L87M | Phosphonate ABC transporter, ATPase | CMM_0376 | G455804->T | S9 |  |
| R435G | Hypothetical protein | CMM_0482 | C561362->G |  |  |
| P224S | GntR-family transcriptional regulator | CMM_0806 | C910375->T |  |  |
| H37R | Transcriptional regulator, HipB/XRE-type | CMM_1284 | A1458166->G |  |  |
| E228G | O-succinylbenzoate-CoA ligase | CMM_0577 | A660294->G | S10 |  |
| E228-> frame shift | Beta-glucosidase | CMM_0082 | GCT122241->G | L1 | **Apoplastic adaptation** |
| A72V | Phosphoribosylamine-glycine ligase | CMM_0754 | G848708->A |  |  |
| G103D | Restriction endonuclease | CMM_2466 | C2781803->T |  |  |
| G103D | Restriction endonuclease | CMM_2466 | C2781803->T | L2 |  |
| A634V | Acyl-CoA oxidase | CMM_0813 | C919435->T | L3 |  |
| G103D | Restriction endonuclease | CMM_2466 | C2781803->T |  |  |
| G103D | Restriction endonuclease | CMM_2466 | C2781803->T | L6 |  |
| ∆A456, ∆T457 | Membrane-bound tyrosin-protein phosphatase | CMM_2386 | GGCGACC2690286->G | L7 |  |
| A26D | NUDIX hydrolase | CMM_1552 | G1758202->T | L8 |  |
| R20H | Hypothetical protein | CMM_2046 | G2309671->A |  |  |
| G196->frame shift | Glutathione S-transferase | CMM_2121 | T2397731->TGCGGCTTC | L10 |  |
