## Supplementary material for "Tissue-Specific Experimental Evolution Reveals Adaptive Trade-Offs in the Plant Vascular Pathogen *Clavibacter michiganensis*": S: Table S1.docx

**Table S1. Sequence substitutions and short indels in vascular and apoplast-adapted clones after 5, 10 and 15 passages**

| **C15** | **C10** | **C5** | **Protein sequence changes** | **Putative annotation** | **Locus tag** | **Mutation** | **Clone** |
| --- | --- | --- | --- | --- | --- | --- | --- |
| Ѵ | Ѵ | Ѵ | synonymous | Phospholipase C | CMM_0504 | G591309->A | S1 |
| Ѵ | Ѵ | Ѵ | R20C | Transcriptional regulator, HipB/XRE-type | **CMM_1284^1^** | C1458114->T^*^ | S1 |
| Ѵ | Ѵ |  | NC | NC | NC^2^ | T790397->C | S1 |
|  |  | Ѵ | NC | NC | NC | C925864->A | S1 |
|  |  | Ѵ | NC | NC | NC | C2928007->CG | S1 |
|  |  | Ѵ | Q300R | Allophanate hydrolase subunit | CMM_0564 | CCA645476->TCG | S1 |
| Ѵ |  |  | NC | NC | NC | G229159->A | S2 |
| Ѵ |  |  | synonymous | Phospholipase C | CMM_0504 | G591309->A | S2 |
|  | Ѵ |  | NC | NC | NC | G988410->C | S2 |
|  |  | Ѵ | S61->SS | Hypothetical protein | CMM_0246 | C308211->CTCG | S2 |
| Ѵ | Ѵ | Ѵ | R20C | Transcriptional regulator, HipB/XRE-type | **CMM_1284** | C1458114->T^*^ | S2 |
|  | Ѵ | Ѵ | NC | NC | NC | A1689137->AG | S2 |
|  | Ѵ |  | NC | NC | NC | C1741610->CG | S2 |
|  | Ѵ |  | NC | NC | NC | CC1881996->GCCCG | S2 |
|  | Ѵ | Ѵ | NC | NC | NC | T2546647->TG | S2 |
|  |  | Ѵ | NC | NC | NC | C127009->G | S3 |
|  |  | Ѵ | NC | NC | NC | C2332120>G | S3 |
|  |  | Ѵ | synonymous | Phospholipase C | CMM_0504 | G591309->A | S3 |
| Ѵ | Ѵ |  | synonymous | Hypothetical protein | CMM_0472 | T548569->C | S3 |
|  | Ѵ |  | NC | NC | NC | T717311->A | S3 |
|  |  | Ѵ | NC | NC | NC | A595018->T | S3 |
|  |  | Ѵ | NC | NC | NC | G925864->A | S3 |
|  |  | Ѵ | NC | NC | NC | GA1576937->TC | S3 |
|  |  | Ѵ | NC | NC | NC | GA1763538->TC | S3 |
|  |  | Ѵ | NC | NC | NC | GA1899616->TC | S3 |
|  |  | Ѵ | NC | NC | NC | AC2546641->GT | S3 |
|  |  | Ѵ | D41->frame shift | Phospho-sugar mutase | CMM_2578 | GGGGAT2915777->CGGGATC | S3 |
| Ѵ |  |  | synonymous | NTP pyrophosphohydrolase | CMM_1199 | C1367906->T | S3 |
| Ѵ | Ѵ |  | H37R | Transcriptional regulator, HipB/XRE-type | **CMM_1284** | A1458166->G^*^ | S3 |
|  | Ѵ | Ѵ | NC | NC | NC | A1689137->AG | S3 |
|  | Ѵ |  | NC | NC | NC | C1741610->CG | S3 |
|  | Ѵ |  | NC | NC | NC | CC1881996->GCCCG | S3 |
| Ѵ | Ѵ |  | L158V | Hypothetical protein | **CMM_2231** | C2524625->G | S3 |
|  | Ѵ |  | NC | NC | NC | T2546647->TG | S3 |
|  |  | Ѵ | NC | NC | NC | AC988419->GT | S4 |
|  | Ѵ |  | synonymous | Membrane protein | CMM_0395 | G474481->A | S4 |
| Ѵ |  |  | A328V | Two component sensor kinase | **CMM_0996** | C1143378->T^*^ | S4 |
| Ѵ | Ѵ |  | H37R | Transcriptional regulator, HipB/XRE-type | **CMM_1284** | A1458166->G | S4 |
| Ѵ |  | Ѵ | NC | NC | NC | A1689137->AG | S4 |
| Ѵ | Ѵ | Ѵ | P68S | Metallo-beta-lactamase | **CMM_1971** | A2227810->G^*^ | S4 |
|  | Ѵ |  | NC | NC | NC | AC2392674->GT | S4 |
|  | Ѵ |  | NC | NC | NC | T2414382->A | S4 |
| Ѵ |  | Ѵ | NC | NC | NC | T2546647->TG | S4 |
|  |  | Ѵ | NC | NC | NC | AC988419->GT | S5 |
|  | Ѵ | Ѵ | NC | NC | NC | AA1458046->TC | S5 |
|  | Ѵ | Ѵ | NC | NC | NC | A1689137->AG | S5 |
|  | Ѵ | Ѵ | NC | NC | NC | T2546647->TG | S5 |
|  |  | Ѵ | NC | NC | NC | GA1899616->TC | S5 |
|  |  | Ѵ | synonymous | Phospholipase C | CMM_0504 | G591309->A | S6 |
|  | ^3^ | Ѵ | NC | NC | NC | C127004->G | S6 |
|  |  | Ѵ | NC | NC | NC | C127009->G | S6 |
| Ѵ |  |  | synonymous | 4-phosphopantetheinyl transferase | CMM_0125 | C185994->T | S6 |
| Ѵ |  |  | NC | NC | NC | TC355914->C | S6 |
|  | Ѵ | Ѵ | NC | NC | NC | G925864->A | S6 |
|  | Ѵ | Ѵ | NC | NC | NC | GA1899616->TC | S6 |
|  | Ѵ | Ѵ | NC | NC | NC | AC2546641->GT | S6 |
|  |  | Ѵ | NC | NC | NC | C233212->A | S6 |
|  |  | Ѵ | NC | NC | NC | GG499078->CC | S6 |
|  |  | Ѵ | NC | NC | NC | AC988419->GT | S6 |
|  |  | Ѵ | NC | NC | NC | C1912409->G | S6 |
|  |  | Ѵ | NC | NC | NC | A2342758->G | S6 |
|  |  | Ѵ | NC | NC | NC | G2371031->A | S6 |
|  |  | Ѵ | NC | NC | NC | T2371036->C | S6 |
|  |  | Ѵ | NC | NC | NC | GC2992290->TC | S6 |
|  |  | Ѵ | NC | NC | NC | C64549->T | S7 |
| Ѵ | Ѵ | Ѵ | A118->AAA | Metal-transporting ATPase | **CMM_1037** | C1187404->CGCTGCT | S7 |
| Ѵ | Ѵ |  | NC | NC | NC | A1689137->AG | S7 |
| Ѵ | Ѵ |  | synonymous | Efflux MFS permease | CMM_1512 | G1713621->A | S7 |
| Ѵ |  |  | NC | NC | NC | C1741610->CG | S7 |
| Ѵ | Ѵ |  | NC | NC | NC | T2546647->TG | S7 |
|  |  | Ѵ | NC | NC | NC | T790397->C | S8 |
|  |  | Ѵ | NC | NC | NC | A1689137->AG | S8 |
|  | Ѵ | Ѵ | T209-> frame shift | Two component sensor kinase | **CMM_0995** | G1142320->GC | S8 |
|  | Ѵ |  | NC | NC | NC | G1458024->A | S8 |
|  | Ѵ | Ѵ | NC | NC | NC | T2546647->TG | S8 |
|  |  | Ѵ | NC | NC | NC | CC1881996->GCCCG | S8 |
|  |  | Ѵ | NC | NC | NC | T790397->C | S9 |
|  |  | Ѵ | NC | NC | NC | C1741463->CG | S9 |
|  |  | Ѵ | NC | NC | NC | T2546647->TG | S9 |
|  | Ѵ |  | NC | NC | NC | C127004>G | S9 |
|  | Ѵ |  | NC | NC | NC | TC355914->T | S9 |
| Ѵ | Ѵ | Ѵ | L87M | Phosphonate ABC transporter, ATPase | **CMM_0376** | G455804->T^*^ | S9 |
| Ѵ | Ѵ | Ѵ | R435G | Hypothetical protein | **CMM_0482** | C561362->G | S9 |
| Ѵ |  |  | P224S | GntR-family transcriptional regulator | **CMM_0806** | C910375->T^*^ | S9 |
| Ѵ | Ѵ | Ѵ | H37R | Transcriptional regulator, HipB/XRE-type | **CMM_1284** | A1458166->G | S9 |
|  | Ѵ |  | NC | NC | NC | GA1899616->TC | S9 |
|  | Ѵ |  | NC | NC | NC | T2414382->A | S9 |
| Ѵ | Ѵ | ^3^ | NC | NC | NC | TC355914->T | S10 |
| Ѵ |  |  | E228G | O-succinylbenzoate-CoA ligase | **CMM_0577** | A660294->G | S10 |
| Ѵ |  |  | E228-> frame shift | Beta-glucosidase | **CMM_0082** | GCT122241->G^*^ | L1 |
| Ѵ | Ѵ |  | A72V | Phosphoribosylamine-glycine ligase | **CMM_0754** | G848708->A^*^ | L1 |
| Ѵ | Ѵ |  | G103D | Restriction endonuclease | **CMM_2466** | C2781803->T | L1 |
|  | Ѵ |  | NC | NC | NC | C2928007->CG | L1 |
|  |  | Ѵ | NC | NC | NC | GG499078->CC | L1 |
|  |  | Ѵ | NC | NC | NC | T790397->C | L1 |
|  |  | Ѵ | NC | NC | NC | T2546647->TG | L1 |
|  |  | Ѵ | NC | NC | NC | A1689137->AG | L1 |
|  |  | Ѵ | T54S | 50S ribosomal protein L4 | CMM_2616 | G2942035->C | L1 |
|  |  | Ѵ | NC | NC | NC | C127004->G | L2 |
| Ѵ |  | Ѵ | NC | NC | NC | C25499->T | L2 |
|  |  | Ѵ | NC | NC | NC | AC988419->GT | L2 |
|  | Ѵ | Ѵ | NC | NC | NC | A1689137->AG | L2 |
| Ѵ |  |  | NC | NC | NC | CC1881996->GCCCG | L2 |
| Ѵ | Ѵ | Ѵ | NC | NC | NC | T2546647->TG | L2 |
| Ѵ | Ѵ | Ѵ | G103D | Restriction endonuclease | **CMM_2466** | C2781803->T^*^ | L2 |
| Ѵ | Ѵ |  | synonymous | Subtilisin-like serine protease | CMM_2536 | G2857977->A | L2 |
| Ѵ |  |  | NC | NC | NC | TC355914->T | L3 |
| Ѵ | Ѵ |  | A634V | Acyl-CoA oxidase | **CMM_0813** | C919435->T^*^ | L3 |
|  | Ѵ |  | P95L | Sugar ABC transporter, ATP-binding protein | **CMM_1244** | C1414665->T | L3 |
|  |  | Ѵ | NC | NC | NC | CC1881996->GCCCG | L3 |
|  | Ѵ | Ѵ | NC | NC | NC | A1689137->AG | L3 |
| Ѵ | Ѵ | Ѵ | NC | NC | NC | T2546647->TG | L3 |
| Ѵ | Ѵ | Ѵ | G103D | Restriction endonuclease | **CMM_2466** | C2781803->T^*^ | L3 |
|  |  | Ѵ | NC | NC | NC | C3278463->T | L3 |
|  |  | Ѵ | G555R | ATP-dependent DNA helicase | CMM_0488 | G567942->A | L4 |
| Ѵ | Ѵ | Ѵ | NC | NC | NC | A1689137->AG | L4 |
|  | Ѵ | Ѵ | NC | NC | NC | T2546647->TG | L4 |
| Ѵ |  |  | NC | NC | NC | C2928007->CGG | L4 |
|  |  | Ѵ | NC | NC | NC | T790397->C | L5 |
|  |  | Ѵ | A383T | tRNA (5-methylaminomethyl-2-thiouridylate)-methyltr | CMM_1403 | G1590720->A | L5 |
| Ѵ |  |  | NC | NC | NC | TC355914->T | L5 |
| Ѵ |  | Ѵ | NC | NC | NC | A1689137->AG | L5 |
|  |  | Ѵ | NC | NC | NC | CC1881996->GCCCG | L5 |
| Ѵ |  | Ѵ | NC | NC | NC | T2546647->TG | L5 |
|  |  | Ѵ | NC | NC | NC | T790397->C | L6 |
| Ѵ | Ѵ | Ѵ | NC | NC | NC | A1689137->AG | L6 |
|  |  | Ѵ | NC | NC | NC | G988410->C | L6 |
|  |  | Ѵ | NC | NC | NC | CC1881996->GCCCG | L6 |
|  |  | Ѵ | NC | NC | NC | T2546647->TG | L6 |
|  |  | Ѵ | NC | NC | NC | T790397->C | L7 |
|  |  | Ѵ | NC | NC | NC | G988410->C | L7 |
|  |  | Ѵ | NC | NC | NC | CC1881996->GCCCG | L7 |
|  | Ѵ |  | NC | NC | NC | G623820->GC | L7 |
| Ѵ | Ѵ | Ѵ | NC | NC | NC | A1689137->AG | L7 |
| Ѵ | Ѵ | Ѵ | NC | NC | NC | T2546647->TG | L7 |
| Ѵ | Ѵ |  | ∆A456, ∆T457 | Membrane-bound tyrosin-protein phosphatase | **CMM_2386** | GGCGACC2690286->G | L7 |
| Ѵ | Ѵ |  | synonymous | Aspartyl-tRNA synthetase | CMM_0561 | G641842->A | L8 |
| Ѵ |  | Ѵ | NC | NC | NC | A1689137->AG | L8 |
| Ѵ | Ѵ |  | A26D | NUDIX hydrolase | **CMM_1552** | G1758202->T | L8 |
| Ѵ |  |  | NC | NC | NC | CC1881996->GCCCG | L8 |
| Ѵ | Ѵ |  | R20H | Hypothetical protein | **CMM_2046** | G2309671->A | L8 |
|  |  | Ѵ | NC | NC | NC | T2546647->TG | L8 |
|  |  | Ѵ | NC | NC | NC | T790397->C | L8 |
|  |  | Ѵ | NC | NC | NC | G925864->A | L9 |
|  |  | Ѵ | NC | NC | NC | AC2546641->GT | L9 |
| Ѵ |  |  | NC | NC | NC | T2546647->TG | L9 |
|  | Ѵ |  | L28->LLAPL | Mlybdopterin biosynthesis protein | **CMM_2314** | C2618298->CGAGCGGGGCGAG | L9 |
|  | Ѵ |  | NC | NC | NC | T790397->C | L10 |
| Ѵ | Ѵ | Ѵ | NC | NC | NC | A1689137->AG | L10 |
| Ѵ | Ѵ |  | G196->frame shift | Glutathione S-transferase | **CMM_2121** | T2397731->TGCGGCTTC^*^ | L10 |
| Ѵ | Ѵ |  | synonymous | RpiR family transcriptional regulator | CMM_2189 | G2484929->A | L10 |
|  |  | Ѵ | NC | NC | NC | T2546647->TG | L10 |
|  |  | Ѵ | D41-> frame shift | Phospho-sugar mutase | **CMM_2578** | GGGGAT2915777->CGGGATC | L10 |

^1^Mutation that resulted in protein sequence alterations are marked in bold

^2^NC – mutation occurred in non-coding areas

^3^Sequencing of S6 from cycle 10 and S10 from cycle 5 did not pass quality control and was removed from the analyses

^*^Mutation was confirmed by Sanger sequencing at C15
