## Supplementary material for "Tissue-Specific Experimental Evolution Reveals Adaptive Trade-Offs in the Plant Vascular Pathogen *Clavibacter michiganensis*": S: Table S2.docx

**Table S2: Bacterial strains used in this study**

| **Strain** | **Relevant characteristic** | **Reference** |
| --- | --- | --- |
| ***Escherichia coli*** | | |
| DH5α | F- ompT hsdSB(rB- mB-) gal dcm (DE3) pRARE, Cm^R^ | Invitrogen (San Diego, CA, USA) |
| ***Clavibacter michiganensis*** | | |
| NCPPB382 | Wild type (WT) parent strain | [1] |
| S1 | Vascular adapted clone collected after 15 passages | This study |
| S2 | Vascular adapted clone collected after 15 passages | This study |
| S3 | Vascular adapted clone collected after 15 passages | This study |
| S4 | Vascular adapted clone collected after 15 passages | This study |
| S5 | Vascular adapted clone collected after 15 passages | This study |
| S6 | Vascular adapted clone collected after 15 passages | This study |
| S7 | Vascular adapted clone collected after 15 passages | This study |
| S8 | Vascular adapted clone collected after 15 passages | This study |
| S9 | Vascular adapted clone collected after 15 passages | This study |
| S10 | Vascular adapted clone collected after 15 passages | This study |
| L1 | Apoplast adapted clone collected after 15 passages | This study |
| L2 | Apoplast adapted clone collected after 15 passages | This study |
| L3 | Apoplast adapted clone collected after 15 passages | This study |
| L4 | Apoplast adapted clone collected after 15 passages | This study |
| L5 | Apoplast adapted clone collected after 15 passages | This study |
| L6 | Apoplast adapted clone collected after 15 passages | This study |
| L7 | Apoplast adapted clone collected after 15 passages | This study |
| L8 | Apoplast adapted clone collected after 15 passages | This study |
| L9 | Apoplast adapted clone collected after 15 passages | This study |
| L10 | Apoplast adapted clone collected after 15 passages | This study |
| LB1 | LB culture adapted clone collected after 15 passages | This study |
| LB2 | LB culture adapted clone collected after 15 passages | This study |
| LB3 | LB culture adapted clone collected after 15 passages | This study |
| LB4 | LB culture adapted clone collected after 15 passages | This study |
| LB5 | LB culture adapted clone collected after 15 passages | This study |
| LB6 | LB culture adapted clone collected after 15 passages | This study |
| LB7 | LB culture adapted clone collected after 15 passages | This study |
| LB8 | LB culture adapted clone collected after 15 passages | This study |
| LB9 | LB culture adapted clone collected after 15 passages | This study |
| LB10 | LB culture adapted clone collected after 15 passages | This study |
| NCPPB382:p*celA:GUS* | NCPPB382 carrying pHN216:p*celA*:*GUS* used for p*celA* promoter activity assay, Neo^R^ | This study |
| S2:p*celA:GUS* | S2 pHN216:p*celA*:*GUS* used for p*celA* promoter activity assay, Neo^R^ | This study |
| S4:p*celA:GUS* | S4 pHN216:p*celA*:*GUS* used for p*celA* promoter activity assay, Neo^R^ | This study |
| S9:p*celA:GUS* | S9 pHN216:p*celA*:*GUS* used for p*celA* promoter activity assay, Neo^R^ | This study |
| NCPPB382:p*gyrB:GUS* | NCPPB382 carrying pHN216:p*gyrB:GUS* used for p*gyrB* promoter activity assay, Neo^R^ | This study |
| S2: p*gyrB:GUS* | S2 carrying pHN216:p*gyrB:GUS* used for p*gyrB* promoter activity assay, Neo^R^ | This study |
| S4: p*gyrB:GUS* | S4 carrying pHN216:p*gyrB:GUS* used for p*gyrB* promoter activity assay, Neo^R^ | This study |
| S9: p*gyrB:GUS* | S9 carrying pHN216:p*gyrB:GUS* used for p*gyrB* promoter activity assay, Neo^R^ | This study |
| S2:pMe*groEL*:*celA*-HA | S2 carrying pCMIAR: pMe*groEL*:*celA*-HA for overexpression of CelA fused to HA tag, Gnt^R^ | This study |
| S4: pMe*groEL*:*celA*-HA | S4 carrying pCMIAR: pMe*groEL*:*celA*-HA for overexpression of CelA fused to HA tag, Gnt^R^ | This study |
| S9: pMe*groEL*:*celA*-HA | S9 carrying pCMIAR: pMe*groEL*:*celA*-HA for overexpression of CelA fused to HA tag, Gnt^R^ | This study |
| NCPPB382:p*CMP1*:*EGFP* | NCPPB382 carrying the EGFP expression plasmid pK2-22, Neo^R^ | [2] |
| S2:p*CMP1*:*EGFP* | S2 carrying the EGFP expression plasmid pK2-22, Neo^R^ | This study |
| S4:p*CMP1*:*EGFP* | S4 carrying the EGFP expression plasmid pK2-22, Neo^R^ | This study |
| NCPPB382Ω1284 | Marker exchange mutant in CMM_1284 in the background of NCPPB382, Neo^R^ | This study |

*Neo^R^ and Gnt^R^ respectively indicate neomycin and gentamicin resistance
