## Supplementary material for "Tissue-Specific Experimental Evolution Reveals Adaptive Trade-Offs in the Plant Vascular Pathogen *Clavibacter michiganensis*": S: Table S3.docx

**Table S3: Plasmids used in this study**

| **Plasmid** | **Relevant characteristic** | **Reference** |
| --- | --- | --- |
| pHN216 | *E. coli-Clavibacter* shuttle vector, based on the replicon of pCM2, Neo^R^/Km^R^, Gnt^R^ | [1] |
| pHN216:p*cel*:*GUS* | pHN216 derivative introduced with *E. coli* *GUS* gene under the control of the Cm *celA* (pCM1_0020*)* promoter. Used for promoter activity assay, Neo^R^/Km^R^ | This study |
| pHN216:p*gyrB*:*GUS* | pHN216 derivative introduced with *E. coli* *GUS* gene under the control of the Cm *gyrB* (CMM_0006*)* promoter. Used for promoter activity assay, Neo^R^/Km^R^ | This study |
| pK2-22 | *E. coli-Clavibacter* shuttle vector, based on the replicon of pCM1, expressing EGFP under the control of the p*CMP1* promoter. Used for in planta localization analyses, Neo^R^/Km^R^ | [2] |
| pMA-RQ:Cmp | Cloning vector for production of Cm integration plasmids carrying the p*CMP1* promoter, MCS and 3×HA tag, Amp^R^ | [3] |
| pCMAT | pMA-RQ:Cmp introduced with ***aac(3)-IV*** upstream to the p*CMP1* promoter, Gnt^R^, Amp^R^ | This study |
| pCMAT:1284 | pCMAT MCS was introduced with a fragment containing the 5' flanking region of CMM_1284-nptII-3' flanking region of CMM_1284. Used for generation of CMM_1284 maker exchange mutant. Neo^R^/Km^R^, Gnt^R^ Amp^R^ | This study |
| pCMIARG | Modified pCMAT variant replacing pCMP1 promoter with Microbacterium esteraromaticum groEL (FVO59_01375) promoter (pMe*groEL*) and with the 89863–90317 region of Cm NPPB382 (AM711867) cloned downstream to the 3×HA tag. Used for Cm overexpression via genomic integration, Gnt^R^, Amp^R^ | This study |
| pCMIARG:*celA* | pCMIARG MCS introduced with Cm *celA* ORF (pCM1_0020) in frame with the C-terminal 3×HA tag. Used for CelA overexpression. Gnt^R^, Amp^R^ | This study |

*Neo^R^, Km^R^, Gnt^R^ and Amp^R^ indicate neomycin, kanamycin, gentamicin and ampicillin resistance, respectively
