## Supplementary material for "Tissue-Specific Experimental Evolution Reveals Adaptive Trade-Offs in the Plant Vascular Pathogen *Clavibacter michiganensis*": S: Table S4.docx

**Table S4. Primers used during this study**

| *Cloning primers* (bold underline represents restriction sites used for cloning) | | | | |
| --- | --- | --- | --- | --- |
| Primer name | Cloned region/gene | Sequence (5’ to 3’) | Destination vector | Generated vector |
| GUScelF | GUS (*uidA*) | GGCTTCCCTACGATCCTTATATGTTACGTCCTGTAGAAACCCC | pHN216 | pHN216:p*cel*:*GUS* |
| GUSgyrF | GUS (*uidA*) | ACGCGGCAGGAGCCACCACTTCATGTTACGTCCTGTAGAAACCCC | pHN216 | pHN216:p*gyrB*:*GUS* |
| GUSR | GUS (*uidA*) | AAA**GAATTC**TCATTGTTTGCCTCCCTGCTG | pHN216 | pHN216:p*cel*:*GUS*/ pHN216:p*gyrB*:*GUS* |
| pcelAF | p*cel* | CCC**AAGCTT**GGCGCTCGGTGCCGACGTCG | pHN216 | pHN216:p*cel*:*GUS* |
| pcelAR | p*cel* | GGGGTTTCTACAGGACGTAACATATAAGGATCGTAGGGAAGCC | pHN216 | pHN216:p*cel*:*GUS* |
| pgyrBF | p*gyrB* | CCC**AAGCTT**CCGCAAGGTGTTCGGCG | pHN216 | pHN216:p*gyrB*:*GUS* |
| pgyrBR | p*gurB* | GGGGTTTCTACAGGACGTAACATGAAGTGGTGGCTCCTGCCGCGT | pHN216 | pHN216:p*gyrB*:*GUS* |
| GntF | ***aac(3)-IV*** | AA**GGTACC**CCCGCCAGCCTCGCAGAG | pMA-RQ:Cmp | pCMAT |
| GntR | ***aac(3)-IV*** | CC**AAGCTT**TCAGCCAATCGACTGGCGAG | pMA-RQ:Cmp | pCMAT |
| 5P1284F | **5' CMM_1284** | AAACTGCA**GGATCC**TCGACGGTGATGC | pCMAT | pCMAT:1284 |
| 5P1284R | **5' CMM_1284** | CTTGCGGCAGCGTGAAGCTTGACGGTGGTGACCATGGCAC | pCMAT | pCMAT:1284 |
| NeoF | ***nptII*** | GTGCCATGGTCACCACCGTCAAGCTTCACGCTGCCGCAAG | pCMAT | pCMAT:1284 |
| NeoR | ***nptII*** | GTGGCGGGTCAGCTTCTGATTGGCAGGTTGGGCGTCGCTTG | pCMAT | pCMAT:1284 |
| 3P1284F | **3' CMM_1284** | CAAGCGACGCCCAACCTGCCAATCAGAAGCTGACCCGCCAC | pCMAT | pCMAT:1284 |
| 3P1284R | **3' CMM_1284** | GGGG**TCTAGA**TCACGGGCGGCTTCGTC | pCMAT | pCMAT:1284 |
| GroELF | **pMe*groEL*** | CCC**AAGCTT**CGCGGCTTGCACTCTCATGGG | pCMAT | pCMIARG |
| GroELR | **pMe*groEL*** | TTT**GGATCC**CATTTCTTCTCGAGCACGACG | pCMAT | pCMIARG |
| IntchpBF | **5' *chpB*** | AA**GAGCTC**CCGGTCGGATGCGGGGATAC | pCMAT | pCMIARG |
| IntchpBR | **5' *chpB*** | CC**GAATTC**GCCGGCGGCCGCGAGTACG | pCMAT | pCMIARG |
| CelAF | ***celA*** | GTGCTCGAGAAGAAATGGGATCCTGCGAAAGGCGTCTGTGCAAG | pCMIARG | pCMIARG:*celA* |
| CelAR | ***celA*** | CTAGACTGCAGCCCGGGGTCGACGTGCACAGGGTACAAGCGGG | pCMIARG | pCMIARG:*celA* |
| *Mutation conformation and Sanger sequencing primers* | | | | |
| Locus tag | **Mutation** | F primer | R primer | |
| CMM_1284 | ***nptII* insertion** | CCTTCATCCTCCGAGCCTA | AAGGAGATCTGATGGAGGCAT | |
| CMM_1284 | **SNP/IDEL** | GTCGGGTGAGGCGTCGGGGTGGCGGGC | CCCGTCACGGTGCGGCCGGCAGAGCGC | |
| CMM_0082 | **SNP/IDEL** | GCGAGACATACGGTGAGGAC | GAGATCGACGTCCATGCCAG | |
| CMM_0376 | **SNP/IDEL** | GGATCCGGGAAGTCGACC | GGATGAGGTCGAGCAGGC | |
| CMM_0754 | **SNP/IDEL** | GATCCTGGTACTCGGTTCCG | GTTGAACTCGATGACCCGGA | |
| CMM_0806 | **SNP/IDEL** | GAGACGGTCCGCAACAAGG | TCAGGCGTCGTGGTCGAC | |
| CMM_0813 | **SNP/IDEL** | TTCAACCGGCACCAGGAC | ACCCCTTCTTCTTCTCGCTC | |
| CMM_0996 | **SNP/IDEL** | ATGGCCGCTAGGCCTCCGG | CCGGTCGCCCCTCTCGCGATC | |
| CMM_1971 | **SNP/IDEL** | CAGTGGGCTAGTGGTCAAGG | CGTCAGAGGCATCGTCATCA | |
| CMM_2121 | **SNP/IDEL** | CAGATCACGCTCGACCTCTC | CTTGAAGTGGCCGTGGTACA | |
| CMM_2466 | **SNP/IDEL** | CCGTCCGTGATCCTGCTC | CTCCATCCCGCTGACCAC | |
| *RT-qPCR primers* | | | | |
| Gene | **Locus tag** | F primer | R primer | |
| *celA* | pCM1_0020 | CATGACTACCCCTCGACCATTTAC | AATGTCCTTCTTCGCCAGGTATC | |
| *aglC* | CMM_2797 | CACGAGATCTACCGCGAGTG | GAGGTAGCTGAAGTTGAACGCC | |
| *chpC* | CMM_0052 | ATCTTCCACAACTGACTTTTTGCT | TGAGATAGCTTGATATACACCATCG | |
| *chpE* | CMM_0039 | GGTACGTGGTCATAGCAAAACA | CGCAACTACGCTTCCTACTTC | |
| *chpG* | CMM_0059 | GCTCAACTCGCCCATCTACTT | CAGTGCAGGGTCATGTTGTC | |
| *xysA* | CMM_1673 | TCATCAACGAGTACAACACCGAC | TGTTCCTCAGCTCTTCGATGTC | |
| *nagA* | CMM_0049 | GGACAGTTGAGCCTCGTATCC | GTACGAGTATGTAAGGAGGTGC | |
| *wzy* | CMM_0715 | GTCTTCCTGCCGTTCCTGATC | GGATGACGATCGAGTAGACCATC | |
| *wcnE* | CMM_0718 | CAAGAACCTGAAGCGCCTCA | TAGCCGGCCAGTTTGATGTC | |
| *wcoA* | CMM_0819 | GGAACGACGTGATGGACAACT | TTGTTCCAGATGTCCACGTGTC | |
| *wcmF* | CMM_1597 | GGAACATCAACCAGAACAGCG | GAAGTGGTTGGTGCCGATGAT | |
| *wcmH* | CMM_1601 | CCGAACTTCACGCACATCAC | CCCTGCTTCATCCCCATGTT | |
| *gyrA* | CMM_0007 | CTGATGAAGCTGCTCGACAT | CGCCAGGATGTGATTGTACTC | |
